# Single protein molecules separation, tracking and counting in ultra-thin silicon channels

**DOI:** 10.1101/2023.11.09.566381

**Authors:** Shilo Ohayon, Liran Taib, Navneet Chandra Verma, Marzia Iarossi, Ivy Bhattacharya, Barak Marom, Diana Huttner, Amit Meller

## Abstract

Emerging single-molecule protein sensing techniques are ushering in a transformative era in biomedical research. Nevertheless, challenges persist in realizing ultra-fast full-size protein sensing, including loss of molecular integrity due to protein fragmentation, biases introduced by antibodies affinity, identification of proteoforms and low throughputs. Here, we introduce a single-molecule method for parallel protein separation and tracking, yielding multi-dimensional molecular properties used for their identification. Proteins are tagged by dual amino-acid specific labels and are electrophoretically separated by their mass/charge in custom-designed silicon nano-channel. This approach allows us to analyze thousands of individual proteins within a few minutes by tracking their motion during the migration. We demonstrate the power of the method by quantifying a cytokine panel for host-response discrimination between viral and bacterial infections. Moreover, we show that two clinically-relevant splice isoforms of VEGF can be accurately quantified from human serum samples. Being non-destructive and compatible with full-length intact proteins, this method opens up new ways for antibody-free single protein molecule quantification.

---

Next-generation and single-molecule DNA sequencing technologies have sharply reduced the cost and time of high-throughput sequencing, consequently transforming our ability to perform whole genome studies and to quantify mRNA expression levels down to single-cells. To date, however, the ability to routinely and directly quantify proteins’ copy numbers and proteoforms has remained an outstanding challenge.^1–3^ This has largely stemmed from the fact that unlike DNAs, proteins cannot be amplified enzymatically and are natively folded into highly complex structures. Furthermore, proteins often appear in several different proteoforms, including sequence variations, splice variants and post-translational modifications.^4^ This diversity of proteoforms is exceptionally significant for our understanding of the molecular basis for cellular functions and is clinically relevant. Presently, most protein identification methods rely on fingerprinting short proteolytically-produced peptides instead of whole proteins, or rely on using multiple antibodies for immuno-assays protein sensing. Despite significant progress,^5–9^ these approaches are fundamentally limited. For example, digested peptides do not always uniquely represent specific proteins or provide accurate quantification of proteoforms.^4^ Immunosorbent assays may lack the multiplexing ability to sense many proteins without an unavoidable bias or address the quantification of proteoforms.^10^

To address these challenges, a new wave of single-molecule sensing technologies has recently emerged.^1^ ^5,11^ Particularly, single-molecule techniques such as fluorosequencing by Edman degradation, Fluorescence Resonance Energy Transfer (FRET), and nanopores have already shown unprecedented progress toward achieving single-molecule protein analysis.^12–19^ At the same time, significant challenges have remained towards achieving unbiased, highly multiplexed and high throughput protein identification particularly using full-length proteins with single-molecule resolution.^20–22^ Separation of proteins by their mass can greatly facilitate their identification, particularly if this can be combined with amino-acid specific chemistries that provide protein-specific information.^23^ Hereby, we introduce a general method for single protein molecule separation, tracking, identification, and quantification based on multi-dimensional molecular feature extraction in subwavelength deep solid-statechannels. The planar geometry of the channels is designed to load and deliver the proteins towards the imaging area by electrophoresis, whilst the chanel depth is optimized to image their motion over time within a high-resolution focal area. To identify full-length proteins, we apply dual-color biochemical conjugation of amino acids residues combined with single protein molecule separation and sensing.

## Ultra-thin, solid state nano-fluidic device for single protein tracking

A key feature of our method is a nano-channel device that enables not only separation of single protein molecules but also *tracking* them *one by one* over time (Figure 1). Surface immobilization of proteins is a powerful method to facilitate single molecule sensing using FRET or other optical methods.^24^ However, protein immobilization does not allow simple protein mass separation. On the other hand, free diffusion of small proteins in solution highly complicates single molecule tracking, as they quickly drift out of focus. To overcome these limitations, we introduce devices featuring ultra-thin solid channels that physically constrain the proteins within a high-resolution focal area of few hundreds of nanometers in the *z*-direction. Furthermore, a fraction of the nano-channel is fitted *in-situ* with a polymeric plug that sufficiently slows down their diffusion or electro-migration (Figure 1a, 1b). This design permits: (i) very fast delivery of thousands of proteins into the device and towards the separation section, and (ii) electrokinetic mass-based separation of proteins coupled to high SNR (signal to noise ratio) single-molecule sensing (Figure 1c, 1d). Moreover, the electrokinetic induced motion of proteins in the polymeric plug enables single-particle *tracking* of the migrating individual proteins. The nonlinear dynamical migration of the proteins in the gradient polymeric matrix separates them by their mass, producing dynamical velocity tracks for each protein, which adds characteristic information for each protein species. Specific labeling of two amino acids, Cysteins (C) and Lysines (K), provides information regarding proteins amino-acids composition, allowing us to precisely classify multiple proteins simultaneously evading reliance on antibodies.

**Figure 1.**
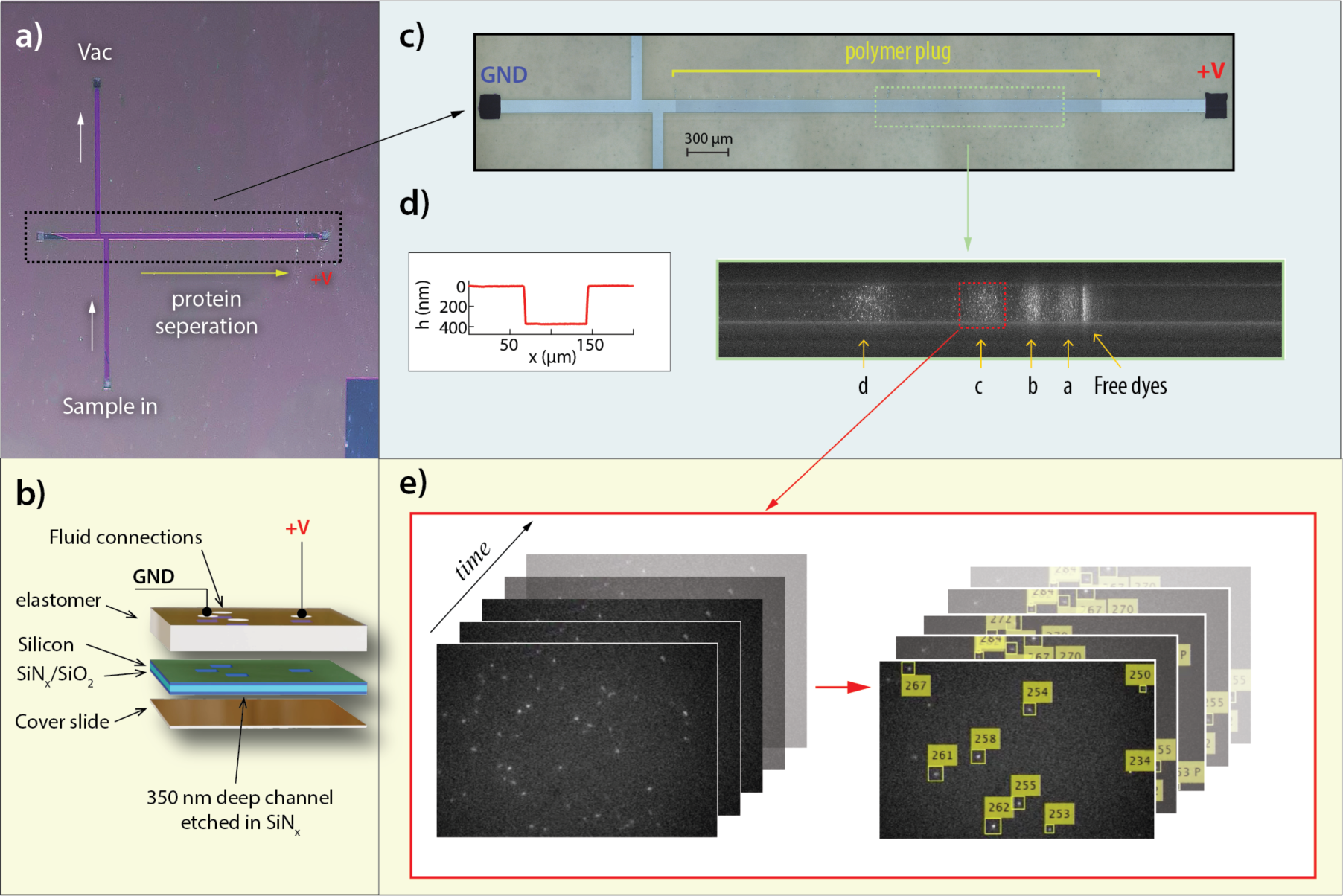
Silicon-based nano-channel for single protein molecule separation imaging and identification. a) A micrograph showing the nanochannel device, viewed from the coverslide side. 350 nm deep channels are etched in the SiNx layer using RIE, and through holes ports are formed using wet etch prior to bonding of glass slide to from a solid channel. b) A schematic side view of the device, showing the glass slide, silicon chip and an elastomer layer bonded at the back of the device to facilitate fluid connections and electrodes placement. c) A magnified white light view of the nano-channel showing the polymer plug section in which proteins are separated by mass/charge. d) Left: A needle profilometer scan of the nanochannel showing a rectangular 350 nm deep channel. Right: Fluorescence Kymograph of a run consisting of 4 labeled proteins (marked a, b, c and d), visibly separated along the channel. e) Single protein molecule trajectories along the channel are enabled by capturing high-resolution videos of a portion of the nanochannel and performing single particle identification and tracking. Alternate lasers (565 nm and 640 nm) excitations yield two independent trajectory plots for the green and red fluorophores (here a single channel is shown).

Devices were fabricated in-house by chemical dry etching selected areas in low-stress silicon-nitride (SiN_x_) layers deposited on silicon wafers (see SI Figure 1), and hermetically sealing the front side of the chip supporting the channels with glass slides, which is anodically bonded to the SiN_x_ layer. The back side of the chip is bonded to a PDMS mold to create fluidic connections (Figure 1b). We found that the solid material fabrication is necessary to support the high width/height aspect ratio of the device, without collapsing.^25^ The filling zone of the devices consists of a double “T” offset junction, which defines a precise sample volume loading of 4 pL. This is an outstanding feature of our device since ultra-small volume analysis represents a significant challenge for the currently established detection methodologies. Specifically, protein samples are loaded into the junction from the entry port by vacuum. At the heart of our devices a 3 mm long by 75 µm wide separation channel with a height of only ∼350 nm (a profilometer scan of the channel is shown in Figure 1d left panel). The device’s internal walls were coated with an anti-stick layer to minimize protein sticking and the effects of an electro-osmotic flow (Methods). The channel is then filled with uncured polyacrylamide and is selectively polymerized *in-situ*, using a UV light illuminated through a microscope objective lens, only at designated “plug” area by mounted the chip on a precision XY motorized stage. Controlling the polymerization location and the UV dose allows us to form a gradient gel density along the channel (Figure 1c), leaving the feed-through sections unpolymerized for fast sample loading. Single-molecule imaging is performed through the glass cover slide in the “imaging zone” (marked by a red rectangle in Figure 1d) located at a distance of roughly 2.5 mm along the channel.

To image the devices, we constructed a custom dual-color single molecule sensing setup that produces uniform illumination of the imaging zone using two laser excitations (532 nm and 640 nm, SI Figure 2). The lasers exposure times were synchronized with an EM-CCD frame acquisition periods and were alternated to allow independent excitation of “green” (Atto 565) and “red” (Atto 643) fluorophores. The emitted light was collected using a high NA microscope objective (Olympus 60x/1.45), split to two channels, filtered and imaged by an EM-CCD camera. The nano-channel device was placed on a piezo-driven *Z* axis stage equipped also with long-travel *X-Y* motors, to allow precise placement of the device at the imaging zone as confirmed using a white light image). To electrokinetically drive the proteins through the nano-channel, a steady voltage was applied using a computer-controlled electrometer. The system was monitored in real-time using a custom LabView program, which in addition to fully control all parts of the system was also used to directly stream the movies to disk from the EM-CCD.

**Figure 2.**
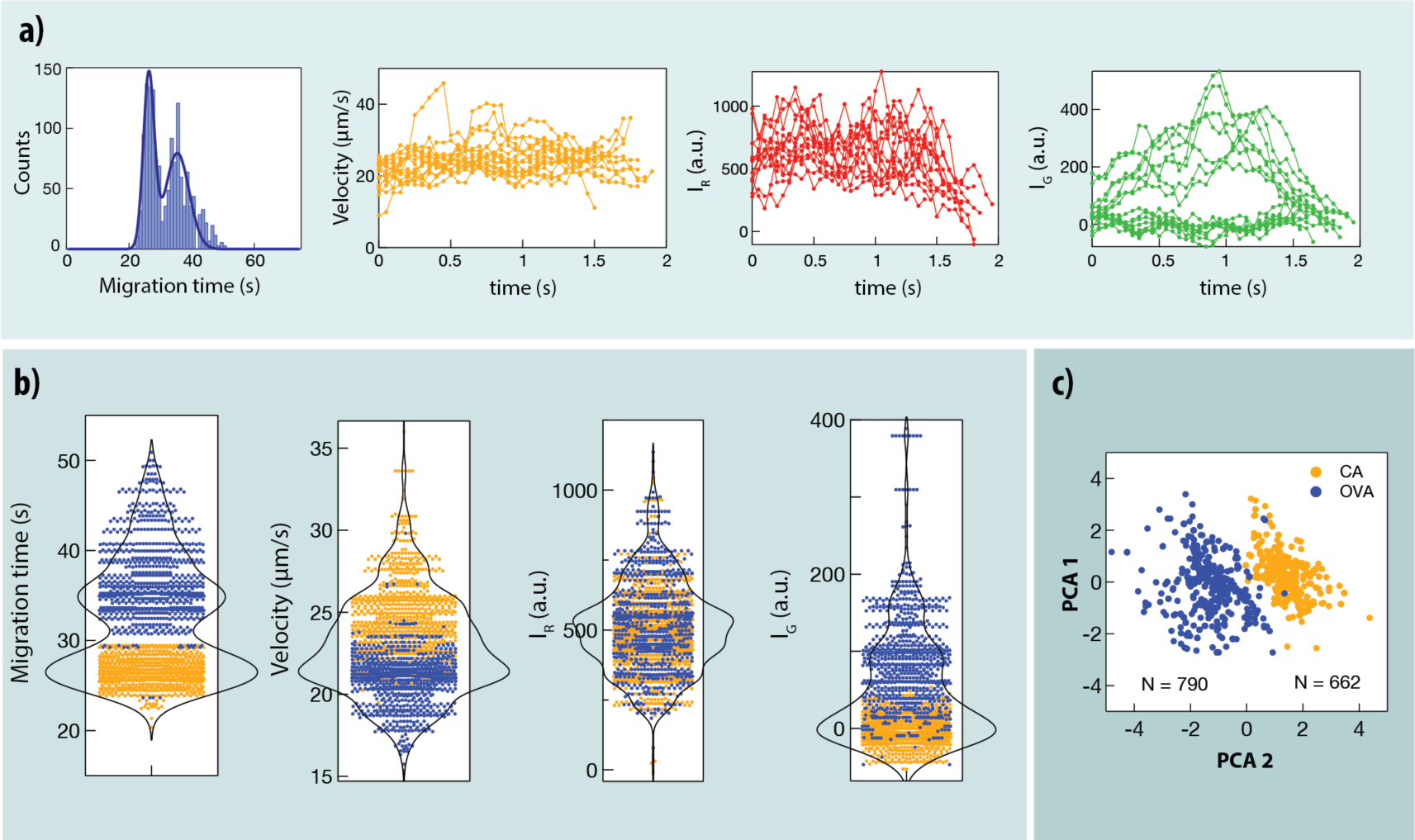
Single protein molecule tracking and identification in nano-channels with dual-color labeling. a) Each protein molecule produces three single molecule trajectories reflecting its in-frame velocity, its fluorescence intensity when excited by the red and by the green lasers. A fourth feature is the protein migration time through the full channel length. b) The mean values of the single molecule tracks (N = 1,452) are plotted as ”violin” graphs, and are subjected to a Gaussian Mixture Model clustering, which annotated two groups. The faster protein (yellow) is identified as the CA (Mw post labeling = 46.78 kDa) and slower protein (blue) as OVA (Mw post labeling = 66.65 kDa), consistent with the fact that CA shows zero green emission (it has no C residues), whereas OVA shows significant green emission. Both clusters show similar red emission, as expected due to similar number of K. c) Principal Component Analysis (PCA) of the 4D information allowed simplified representation in a 2D graph, clearly showing two distinct clusters, as annotated by GMM.

The low profile of the nano-channels in the vertical direction ensures that the proteins remain in focus during their migration, as shown in *Supplementary Movie 1*. We find that this and the high SNR in the single-molecule imaging are essential features for the development of a single-particle tracking algorithm, used to numerate each protein migrating through the camera frame (Figure 1d). To produce continuous particle trajectories, we first localize the particles in each frame and then apply a routine based on Kalman filter that connects the most likely trajectories throughout the entire frame stack (Methods).^26,27^ If the fluorphore intensity fell below a noise threshold at a certain movie frame and re-apear in the next frame our algorithm interpolates the expected location of these particles to create continuos single-molecule dynamic trajectories.

## Amino-acids specific labeling for 4D protein discrimination

To further enhance single protein molecule discrimination beyond their mass/charge ratio, our method quantifies independently the number of two amino-acids, Lysines (K) and cysteins (C) residues. To that end, we modified common chemical conjugation procedures to achieve a higher degree of labeling (DoL: the fraction of K or C labelled per available residues per protein). Proteins were first reduced by TCEP and then denatured by heat in the presence of surfactant sodium dodecyl sulfate (SDS). In most cases we reach 100% DoL for both dyes, and in all cases the DoL > 60% (SI Table 2), as confirmed by UV-Vis and bulk gel (Supporting Information). To validate our analysis tools and demonstrate the method’s capcity to extract 4D information, we prepared mixtures of two dually labeled proteins with easily distinguishable mass difference: Carbonic anhydrase - CA (30 kDa), and Ovalbumin - OVA (44.2 kDa). These proteins harbor similar numbers of K residues (18 and 20, respectively) but significantly different numbers of C residues (0 and 6, respectively, Table 1). A difference of roughly 20 kDa in M_w_ post-labeling is easily discernable in bulk SDS-PAGE (SI Figure 3). Prior to loading of samples to the chip in the designed port, the sample is typically diluted by roughly 10^4^ – 10^8^ fold depending on the original sample concentration, to a typical in-channel concentration of 1 to 100 pM for single-molecule sensing. A voltage is applied between the two platinum electrodes (GND and “+V”) causing proteins to quickly stack at the buffer/gel interface and begin their slow migration in the polymerized section of the nano-channel.

**Figure 3.**
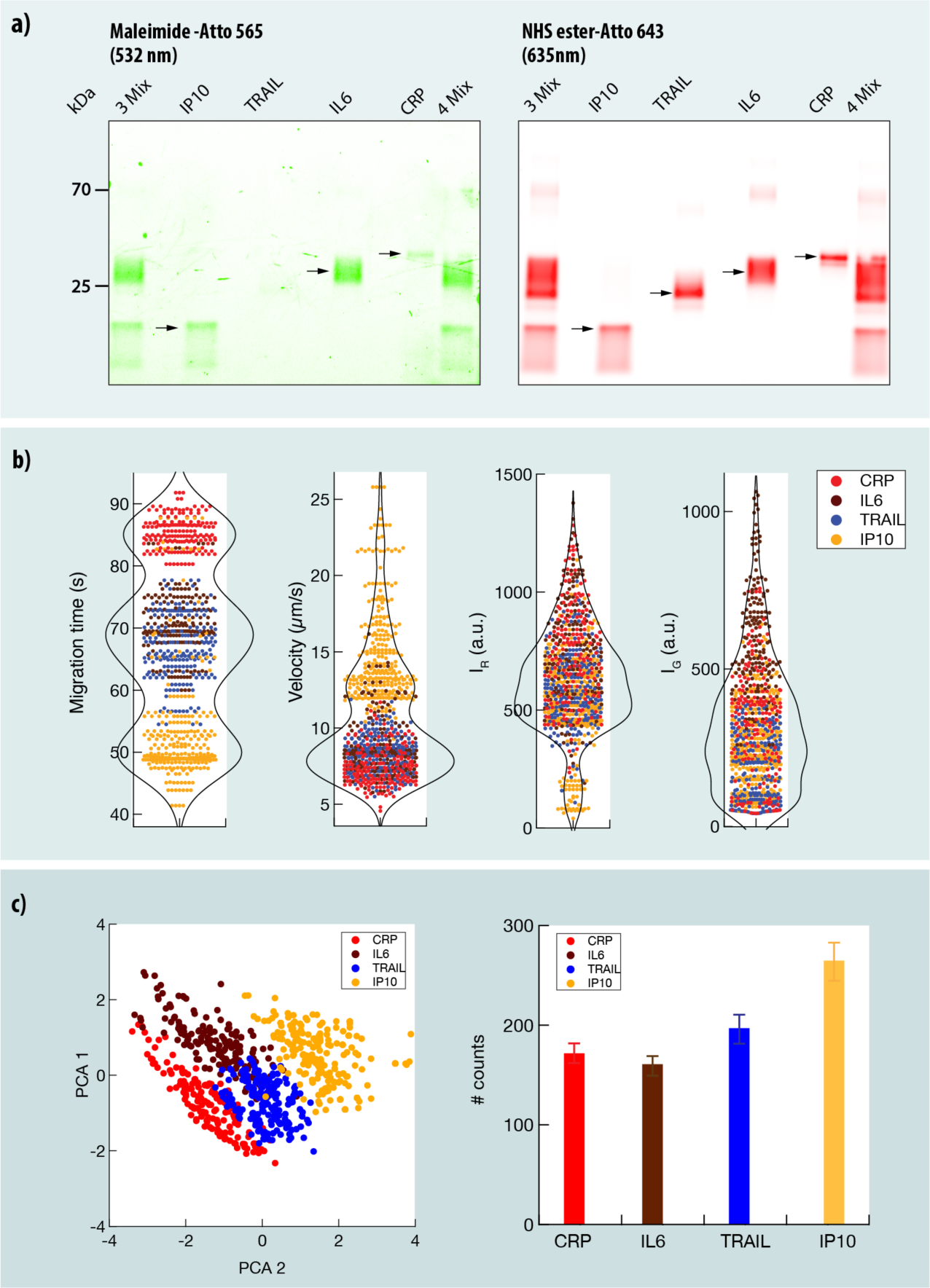
Single protein molecule quantification of a 4 cytokine panel using SPM-trac. a) SDS-PAGE image of dual-color labeled cytokines panel (IP10, Trail, IL6, CRP). All proteins were labelled as described in Methods to >70% yield. b) Violin plots of the 4 cytokine mix run in nano-channels. Despite the similarity in Mw among some of the proteins, SPM-track can distinguish among the 4 cytokines. c) Left: PCA plot with cytokine annotation based on the GMM analysis. The counts of each of the 4 proteins are shown on the right, permitting a direct quantification of the sample.

**Table 1:** In the top panel proteins properties used for validation of the method and for discrimination and quantification of the VEGF isoforms. In the bottom panel properties of the four cytokines, used for hostresponse analysis. The molecular weight post labeling is estimated based on full labelling approximation.

| Protein | M<sub>w</sub> (kDa) | Cysteines | Lysines | M<sub>w</sub> post labeling |
| --- | --- | --- | --- | --- |
| CA | 30 | 0 | 18 | 46.78 |
| OVA | 42 | 6 | 20 | 66.65 |
| VEGF 121 | 14.06 | 9 | 7 | 26.29 |
| VEGF 165 | 19.1 | 16 | 11 | 39.49 |

**Table 1:** In the top panel proteins properties used for validation of the method and for discrimination and quantification of the VEGF isoforms.
| Cytokine | M<sub>w</sub> (kDa) | Cysteines | Lysines | M<sub>w</sub> post labeling |
| --- | --- | --- | --- | --- |
| IP-10 | 8.65 | 4 | 10 | 20.51 |
| TRAIL | 19.6 | 1 | 11 | 30.49 |
| IL-6 | 20.98 | 4 | 14 | 36.57 |
| CRP | 23.2 | 2 | 13 | 36.59 |

As shown in *Supplementary Movie 1* two discernable populations of single molecules particles appear. The particle trajectories were used to produce 4 graphs (Figure 2a): First, the histogram of the individual proteins’ migration times from the beginning of the nano-channel gel plug up to the observation zone (“migration time”). The other three graphs depict single protein tracking over time, namely: (i) the proteins’ in-frame velocities, (ii) the proteins’ fluorescence intensities with red laser excitation (*I*_*R*_), (iii) the proteins’ fluorescence intensities with green laser excitation (*I*_*G*_). For clarity, in Figure 2a we show only a dozen of representative single protein tracks out of roughly 1,452 proteins collected in about 60 s experiment. Notably, the proteins’ the *in-frame* mean velocity is not constant along the channel and depends on their location (〈*v*〉_*frame*_ = *f*(*x*)), where *x* is a coordinate along the channel. The migration time (*t*_*m*_) is a sum of two main terms, the entry time of proteins into the gel plug, and their in-gel migration time both exhibiting nonlinear dependency on M_w_. This nonlinearity is further magnified by applying a gel *density gradient* during the UV polymerization. Consequently, the migration time *t*_*m*_and the in-frame proteins’ velocities 〈*v*〉_*frame*_ exhibit a different dependency on the proteins’, mass/charge ratio that can be used to further enhance the mass separation power.

To analyze the results the mean values of the 4 graphs in Figure 2a accumulated for 1,452 particles, were plotted as “violin plots” histograms (Figure 2b). Each spot on these plots represents the mean values of a single protein trajectory along the channel. In principle, it is possible to fit these distributions *independently* to a mathematical model representing the two species. However, this would not take full advantage of the fact that the information is obtained from a single experiment. Inspired by previous literature,^28^ we treat the whole data using a Gaussian Mixture Model (GMM), which “tags” each particle to its most likely cluster, based on global minimization fit in all 4 graphs. The global GMM analysis annotated each particle as either CA (yellow) or OVA (blue). Starting from the right panel, we see two clear peaks, where the faster proteins (arrival time of roughly 28 s) are denoted as the CA and the heavier proteins (OVA) arrival time are roughly 35-40 s. Consistently, at the green laser excitation the near zero proteins are the CA (yellow) which have *no* C residues and the rest are OVA. The red laser excitation panel is randomly mixed, consistent with the fact that the two proteins have similar number of K residues (18 and 20, respectively). Finally, the GMM correctly annotated the faster proteins on the Velocity plot as the CA (yellow) with mean speeds around 25 µm/s or more, and the slower proteins as the OVA (blue) velocities around 22 µm/s or less. A 2D Principal Component plot of the four-dimensional information is reported in Figure 2c, showing clear separation of the data into two distinct clusters of events. We can use this classification to numerically count the two proteins with high degree of confidence. Figure 2c shows a very small number of misidentified particles (< 0.4 %).To validate the two-cluster classification of the GMM analysis we performed Calinski Harabasz statistical analysis using our 4D tracking data. Our results (SI Figure 8) confirm that two clusters are statistically most probable attributing high confidence to our analysis method.

## Single protein molecule cytokines panel discrimination

Next, we used our method to digitally quantify the relative numbers of a set of cytokines of biomedical relevance. Discrimination between bacterial and viral infections in young patients is often challenging due to the similarity of their clinical symptoms, but is crucial to avoid excess use of antibiotics and prolonging recovery time due to inappropriate treatments.^29^ Diagnostic tests, which are routinely carried out in the clinic, include culture, serology, and nucleic acid-based test (such as RT-PCR), may assist clinicians in determining the source of infection and subsequently the most accurate treatment, but are all directed towards pathogen identification.^30^ An alternative or complementary diagnostic strategy is to analyze the host’s immune response to the infection.^31^ This strategy bypasses the need to determine whether an identified pathogen is the direct cause of the infection or an unrelated colonizer as well as potentially provides better diagnostic outcomes in the case of more complicated cases, such as mixed infections with both virus and bacteria. Circulating host biomarkers can be monitored by enzyme-linked immunosorbent-assay (ELISA) at the point-of-care, however accurate and unbiased quantification remains to be a challenge. Interleukin 6 (IL-6) and C-reactive protein (CRP) are such biomarkers, up-regulated in bacterial infections, which are often used to support pathogen-based diagnosis in the clinic. Large-scale proteomics screen performed recently introduced a novel host-induced viral biomarker, TNF-related apoptosis-inducing ligand (TRAIL).^32^ Additionally IP-10 is a smaller cytokine, which showed elevated levels in both bacterial and viral infections, with a more pronounced increase in the latter case. The combined signature of CRP, IP-10, and TRAIL cytokines resulted in an accurate and robust differential diagnosis of acute bacterial or viral infections, validated in subsequent clinical studies,^33–35^ prompted us to explore the possibility of *single molecule and antibody-free* discrimination among CRP, IP-10, TRAIL, and IL-6 cytokines while quantifying their relative abundance.

First, we optimized our system by using only three cytokines, which were much easier to identify: IL6 and TRAIL have very similar M_w_, as reported in Table 1, but are strikingly different in their number of C (4 and 1, respectively), whereas IP10 is much lighter protein (8.6/20.5 kDa pre/post labeling) and harbors 4 C like IL6. In contrast, both IP10 and Trail harbor a similar number of K (10 and 11, respectively), whereas IL6 harbors 1. SI Figure 12 summarizes the nano-channel measurement using IL6, IP10 and Trail. As expected, the lightest protein IP10 (orange markers) separates readily in both velocity and migration time, whereas IL6 and TRAIL (brown and blue markers, respectively) display markedly different green signal. GMM analysis of the entire data set was used to identify and count the each of the three cytokines. Next, we performed experiments with all four cytokines in a mixture, shown in Figure 3. The fourth cytokine (CRP) molecular weight is 23/37 KDa (pre/post labeling) and it harbors 2 C and 13 K, which should allow it to be discriminated primarily by the red fluorescence intensity. Bulk SDS-PAGE analysis of each protein after dual labelling along a molecular weight ruler is shown in panel a (left-hand panel shows the green excitation and the right-hand panel the red excitation). Using the molecular weight calibration, the gel suggests that the proteins are labelled as expected (see SI Table 2). This was further confirmed by separately analyzing each of the proteins in the UV-Vis spectrometer for CRP and TRAIL (SI Figure 5) and for IP10 and IL6 by migration shift estimation post C labeling (SI Figure 5). The right-most lane in the gel shows all four cytokines in a mixture. Notably, despite the very high quantity of the proteins used in this bulk analysis, a clear identification of the 4 cytokines in bulk is unpractical.

We performed a single-molecule protein analysis of >1,000 proteins in the nano-channel. As before, single particle tracking was performed leading to 4 violin plots (Figure 3b). Global GMM classification annotated the proteins as graphically displayed in the PCA plot (Figure 3c, left panel). Going back to Figure 3b, we tagged the proteins based on the GMM classification: from the in-frame velocity and the migration time graphs, we can easily identify the lighter protein IP10 marked in yellow, as well as the heaviest protein CRP, marked in red. The green intensity is extremely useful in separating TRAIL with its single C (blue) from the other two proteins harboring 4 C, but not from CRP containing 2 C (red). IL6 shows both high green and high red intensities (in brown), and TRAIL (in blue) exhibits relatively low green and low-medium red intensities, as expected. The GMM classification directly counts the number of each of the proteins in the data set, as presented in the right-hand panel of Figure 3c. Hence, using our method, we can potentially offer significantly enhanced sensitivity by a several orders of magnitude to accurately differentiate between bacterial and viral infections, without reliance on antibodies during the sensing/quantification stage.

## Single-molecule quantification of VEGF isoforms

Our method can resolve and quantify small differences in the proteins M_w_ as well as their C and K amino-acid composition. Consequently, it can be applied to resolve and count full-size proteoforms that are not easily distinguished by MS or immunosorbent methods. To demonstrate this strength, we analyze two closely related isoforms of the Vascular Endothelial Growth Factor-A (VEGF-A) protein,^3637^ which rise from alternative splicing of the VEGFA gene.^38^ The ratio between two isoforms, VEGF121 and VEGF165, which is relevant in various cancerous processes^39^ was quantified either when spiked into human serum or endogenously using our method. VEGF has been associated with multiple physiological and pathological processes ranging from vasculogenesis and angiogenesis to vascular disease in the retina and cancer.^40^ In addition to being subjected to proteolytic regulation, VEGF expression is diversified using alternative splicing.^41^ To date, most quantitative studies on VEGF isoforms relied on mRNA expression,^42,43^ however mRNA expression levels do not always correlate with the actual protein content. Notably, the discrimination among VEGF proteoforms is often complicated by the lack of specific antibodies to allow an unbiased quantification of the various types,^36,37,44^ particularly in physiological samples, in which VEGF resides in small amounts. Yet there is increasing evidence that different VEGF isoforms play different physiological roles, often via isoform selective co-receptors.^41^ The VEGF121 and VEGF165 isoforms were also reported to bind VEGF receptors with different affinities^45^ and to play different roles in different types of malignancies.^43,46,47,48^

To show that our method can provide biologically and clinically useful insights from a clinical sample, we targeted VEGF121 and VEGF165 in buffer, in spiked serum sample and endogenously in human serum. To test the quantification accuracy of our method, we first individually labelled VEGF165 and VEGF121 using both Maleimide and NHS ester reactive dyes and analyzed three different stoichiometric mixtures of the two isoforms with *x* = *C*_165_/(*C*_165_ + *C*_121_) using our method *(Supplementary Movie 2)*. Our results, shown in Figure 4a, suggest that the two VEGF isoforms are readily separated based the 4D data (SI Figure 10). We clearly observe correlation between the fast-moving proteins and lower red intensities. This is expected given that VEGF 121 harbors smaller number of K residues as compared with VEGF165 (Table 1). Discrimination in the green channel is not observed, possibly since most of the Atto565 emission is transferred to Atto643 dye via FRET and the two proteins exhibit undistinguishable FRET values. As before, GMM clustering of the results provides a tool to mark and identify the two VEGF isoforms from the single molecule tracking information. We used the isoforms’ clustering results to calculate the in-channel measured ratio *x*_*Channel*_ as a function of the prepared mixture ratio *x_Sample_* as shown in the right panel of Figure 4a. The results show a strong quantitative correlation with a slope of near unity (0.996 ± 0.058), proving that our method can quantify the VEGF isoform with high accuracy and low bias.

**Figure 4.**
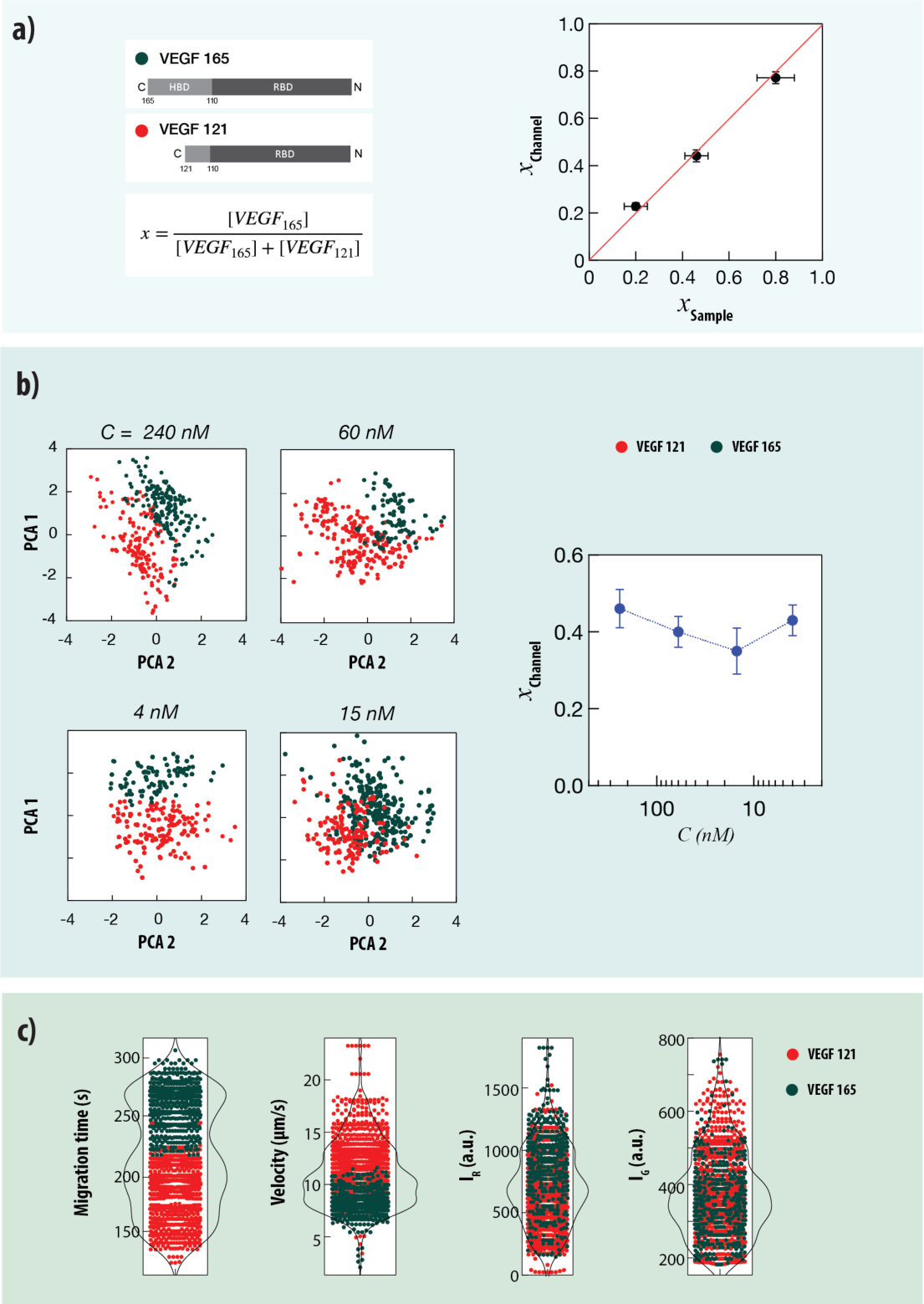
VEGF isoforms separation and digital quantification in buffer, spiked in human serum and endogenously detected from serum. a) Three mixtures of dually labelled VEGF121 and VEGF165 with different relative concentrations *x* = *C*_165_/(*C*_165_ + *C*_121_) analyzed using nano-channels. The 4D information (SI Figure 10) is used to cluster the data using GMM and calculate the ratio of two isoforms. Right: Comparison of the prepared VEGF isoform samples mixture ratio with the single-molecule counting results. A linear regression fit yield a slop of 0.996±0.058 suggesting that the single-molecule counting analysis of the VEGF isoform is quantitative. b) Spike-in of human sera samples with mixture of the VEGF isoforms at 4 different initial concentrations (from 240 nM down to 4 nM) having the same sample ratio × = 0.4. The resulting PCA plots are shown (the 4D information is provided in SI Figure 11). Right: The resulting single-molecule isoform quantification pulled down from the spiked human sera showing the recovery of the isoform ratio as expected *x* = 0.40±0.02. c) Single molecule quantification of endogenous VEGF isoforms from human serum, yielding ratio *x* =0.43±0.02 (n = 1,107). Violin plots results of the 4D tracking analysis are shown.

Given that our method can measure the stoichiometric ratio of the VEGF isoforms, we next challenged its ability to do so starting with a clinical sample of spiked human sera. To that end we spiked sera with VEGF isoform mixtures at four different total concentrations (from 240 nM down to 4 nM) and measured the recovered ratio. The spiked sera samples were first depleted of the most abundant proteins using an affinity column (“High-Select Top14”). Then we recovered the two VEGF isoforms using magnetic beads conjugated in house to a single antibody (Methods). Results for four different total VEGF isoforms concentrations prepared in sera at a ratio *x* = 0.4, are shown in Figure 4b. Discrimination among the two recovered isoforms is straightforward using the 4D data analysis (SI Figure 11). In each case about 10^3^ single molecule trajectories were collected within ∼60 seconds yielding a measured ratio *x* = 0.40 ± 0.02 in good agreement with the spiked value.

Finally, we used our method to detect endogenous VEGF isoforms from the human sera. The workflow for the preparation of the sample is similar to the one developed for the set of spike and recovery VEGF experiments (but obviously no spike-in). The summary of our results is shown in Figure 4c, where about 1,100 single molecule tracks were collected within a few minutes. As before, the separation between VEGF121 and VEGF165 groups in terms of arrival time, average velocity and red emission were clear leading to straightforward GMM based clustering. From these results we measured the ratio of VEGF121 to VEGF165 to be 1.33:1 (x= 0.43 ± 0.02) in the female human serum sample used. These results show that our method is capable of the identification of proteins in low concentrations from clinical samples relevant for diagnostic challenges.

## Discussion

The development of methods for full-size protein identification capable of rapid, multiplexed and high-sensitivity detection is on great demand but still facing challenges. Emerging immunosorbent techniques offer single-molecule sensitivity but suffer from quantification biases when used for detection of many proteins simultaneously, and limited proteoform discrimination. Here we combine molecular weight separation of full-length individual proteins, with molecular quantification of the number of specific amino-acids to precisely identify protein molecules within a complicated protein mix. Four main aspects are the pillars of our method: (1) Crafting nano-channel devices through lithographic methodologies and selectively equipping them with gradient polymer plugs. (2) Near complete dual color labeling of proteins targeting K and C residues. (3) The acquisition of dynamic protein movies, staged using a custom-built experimental setup. (4) Custom-made computer programs to cluster proteins based on 4-D information obtained from the single particle tracking of each protein molecule in the channel. A key feature of the device lies in its ability to slow down the protein’s movement and introducing an element of nonlinearity to their motion, which is driven by voltage inside the polymer plug. As a result, we were able to measure dynamical trajectories over time of single protein molecules, used to uniquely identify proteins of similar mass and discriminate between them even in complex biological media. This approach offers several advantages compared to other single-molecule protein analysis based on nanopores, as we have shown analysis and discrimination of full-length proteins and the parallelization of the readout by tracking the motion of thousands of proteins within a few minutes.

To illustrate the methods’ power we demonstrated a quantitative discrimination among two *VEGF isoforms* in recombinant mixtures, spiked human sera samples with extremely low sample volume and concentrations, and endogenous VEGF proteins. Unlike the classical “sandwich” ELISA assay, our method evades the need for reporter antibodies, while still permitting single-molecule sensing with high throughput. Since our method involves only chemical amino-acid specific conjugation, it is less susceptible to quantification biases and lend itself towards greater multiplexing ability. New conjugation chemistries targeting additional amino-acid residues^1^ or even proteins PTMs^49^ can further enhance the method’s scope to address an extremely broad range of biological questions with single-molecule precision. To further demonstrate the method mutiplexibility we sensed and quantified a cytokine panel, relevant for differentiation between viral and bacterial infections.^32^ Notably, this method can be integrated to enhance whole proteome screening and PTM mapping prior to literally any other single molecule sensing technique, including nanopore based protein sequencing, sm-FRET based protein recognition, fluorosequencing, as well as other emerging approaches involving N-terminal binders, or even future MS profiling.^12^

## Methods

### Single particle tracking algorithm

We acquired video-clips of the imaging zone using custom LabView code that synchronizes the alternate dual laser excitation with the EM-CCD (Andor iXon) exposure. The clips are 512 × 512 pixels in size at a frame rate of 20 fps. Each protein can be seen as a bright particle moving across the channel. To achieve single-particle tracking we first determine the proteins’ location and their area size (number of pixels it is spread on) in each frame. The images are corrected for background based on a mask consisting of the mean value of each pixel from the first 100 frames in the experiment, in which no proteins are present. Then we apply a Laplacian of Gaussian filter on the image in order to find the border lines of the objects in the image. With this information the images are segmented creating a binary mask representing the particles in the image. This allows counting the number of objects (proteins) in the image, finding their area size, and calculating their intensity and center of gravity. The center of gravity corresponds to the protein location in the image and is calculated using:

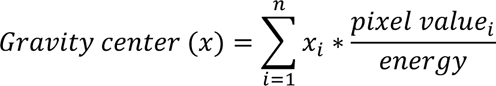

The *x* index of the center of gravity is calculated by the sum of the × index of each pixel (i) from the protein segment multiplied by the photon count of that pixel divided by the total energy of the protein segment. The same equation is used to also calculate the *y* index of the center of gravity.

The mean pixel value of 5 × 5 pixels around the protein center of gravity calculates the intensity of the protein. At the end of this step, for each image in the experiment, we achieve a list of all detected proteins with their location, size area, and intensity to be used for the next tracking step.

Next, the video clips are divided into batches of 120 frames each and an overlap of 6 frames between each batch. The location and size of the proteins in a batch are the input for the tracking algorithm. In this multi-object tracking algorithm, we used Kalman filter to attribute each location to a certain protein. The algorithm collects the initial locations of the proteins and by assuming that an object moves in accordance with a particular motion model (such as constant velocity or constant acceleration), the Kalman filter forecasts the protein’s next location. The process noise and measurement noise are taken into account by the filter. Here, the process noise is the difference between the protein’s real motion and its motion model. Similarly, the detection error is referred to as the measurement noise using this algorithm, from which the trajectory of each single protein is defined. Further individual protein trajectory is monitored and if the protein is not detected for a few frames numbers (a threshold frame number), it will predict its location using the Kalman filter until it appears again. In case the protein does not appear again, the algorithm will discard this proteins trajectory. The advantage of Kalman filter is that it can predict the location of a protein even when its fluorescence emission is too weak to be detected in each frame. The final output is the trajectory of many proteins in a batch. In a single batch, it is possible to have a very high number of objects with similar properties, hence complicating accurate tracking. To overcome this issue, we filtered unlikely trajectories, in which a protein stepped against the applied electrical field. From the proteins tracking we extract the three single particle trajectories: (1) in-frame velocity, (2) fluorescence intensities when excited by red laser (640 nm), (3) fluorescence intensities when excited by green laser (532 nm). We find that all proteins (except CA) exhibited significant FRET when excited by the green laser.

### Data clustering and classification

Initial estimation of the classification values was based on K-means using the expected number of clusters. With these initial values we performed Gaussian Mixture Model (GMM) classification which provides: i) The mean value of each parameter per cluster, ii) the covariance matrix of the parameters and clusters, and iii) proportion of each gaussian distribution. Each 4D data point can then be assigned probability value for a certain cluster based on its cartesian proximity to the gaussian distribution. Principle component analysis (PCA) is used only to display the 4D data in the 2 main coordinates, when each data point is being colored based on the cluster it belongs to.

## Materials

EDTA-treated sera samples (NEGSMPL-P-100) were obtained from healthy subjects from RayBiotech. High-Select™ Top14 Abundant Protein Depletion Resins (catalog A36370) and Dynabeads MyOne Tosylactivated (catalog 65501) were purchased from Thermo Fisher Scientific. Mouse IgG kappa (clone MG1-45) and Mouse anti-VEGF antibody (clone L308D10) were purchased from Biolegends. Atto565 maleimide (AD 565-45) and atto643 NHS ester (AD 643-35) were purchased from Atto-tech GmbH. DMSO was purchased from (Sigma). TCEP was purchased from SIGMA. The details of all proteins used in this study are provided in SI Table 1. VivaSpin concentrators with a 30kDa cutoff were purchased from Cytivia. Pre-cast 4-12% Bis-Tris SDS-PAGE were purchased from Thermo Fisher Scientific. 10% SDS was purchased from Biorad.

### Dual-color labeling of proteins

Proteins were resuspended in Maleimide labeling buffer consisting of 10 mM sodium phosophate, 150 mM NaCl and 1 mM SDS (pH 7.4). Disulfide bonds were reduced using 0.5 mM TCEP at 37°C for 30 minutes and denatured at 95°C for 5 minutes. After cooling down at RT for 15 minutes, atto565 maleimide, dissolved in DMSO or in buffer, was added to the protein samples at a ratio of 2.6x fold per C residue and incubated at 25°C for overnight with shaking at 300 rpm. Subsequently, the samples were dialysed against 10 mM sodium phosophate, 150 mM NaCl and 1 mM SDS (pH 8.4, adjusted with 0.2M sodium bicarbonate (pH 9)). Atto643 NHS ester, dissolved in DMSO or in buffer, was added to the protein samples at a ratio of 20x fold per K residue and incubated at 25°C for 1h inside a heat block with shaking at 300 rpm. The labeled samples were diluted 10x fold with 10 mM sodium phosophate and 1 mM SDS (pH 8.4) buffer to adjust the NaCl concentration to 15 mM. Unconjugated fluorophores were removed using at least six buffer exchange washes with the protein concentrator VivaSpin of a 30 kDa cutoff. The final volume of the labeled proteins was adjusted with NHS ester labeling buffer containing 10 mM sodium phosophate, 15 mM NaCl and 1 mM SDS (pH 8.4). Detailed explanation on how the labeling yield was estimated / calculated is provided in the SI. Qualitative analysis for the labeled proteins was carried out by separating a small fraction of the dually labeled protein samples on a 4-12% Bis-Tris SDS-PAGE and imaged using the Pharos scanner (Biorad) with laser excitation at either 532 or 635 nm.

### Conjugation of BSA or antibodies to tosylactivated dynabeads

40 μg of either mouse IgG or mouse anti-VEGF antibody were conjugated to 1 mg of tosylactivated beads according to the manufacturer protocol for 16h at 37°C on a tube rotator. The blocking step was performed in 1X PBS with 0.05% Tween-20 and 0.5% BSA for overnight at 37°C on a tube rotator. Subsequently, the beads were washed three times in 1X PBS with 0.05% Tween-20. Finally, the beads were resuspended at 2.5 mg/ml final concentration using 1X PBS containing 0.05% Tween-20 and 0.02% sodium azide and kept at 4°C for further usage. Similarly, 2 mg of BSA were conjugated to 1 mg of tosylactivated beads according to the manufacturer protocol to obtain BSA-conjugated tosylactivated beads.

### Immunoprecipitation of VEGF165 and VEGF121 isoforms from human serum

Human sera samples from healthy subjects were spiked with different initial concentrations of VEGF121 and VEGF165. These samples were subjected to high abundant protein depletion using the Thermo Scientific™ High-Select™ Top14 Abundant Protein Depletion Resin according to the manufacturer protocol. The diluted high abundant proteins-depleted plasma was adjusted to contain a final concentration of 1X phosphate buffer saline (PBS) supplemented with 10% glycerol and 0.05% Tween-20 and was subjected to centrifugation at 14,000 g at 4°C for 10 minutes. The resulting supernatant was incubated with BSA-conjugated dynabeads (Thermo Fisher) for 1 h at 4°C on a tube rotator. Total protein of the Top14-depleted plasma sample was quantified using the BCA reagent assay as compared to a BSA standard curve. Typical obtained protein concentrations were ∼0.15 mg/ml. After this pre-clearing step, the BSA-conjugated dynabeads were collected on a magnet, the supernatant was transferred into a new tube, and subjected to centrifugation at 10,000 g at 4°C for 2 minutes. Subsequently, 10 mg of total protein of the this cleared sera supernatant was used per immunoprecipitation experiment. The samples were incubated with 2-4 μg of either mouse IgG or mouse anti-VEGF antibody-conjugated to tosylactivated dynabeads (prepared at 0.04 mg antibody per mg beads) at 4°C for overnight on a tube rotator. Anti-VEGF antibody-conjugated beads were collected on a magnet for 2 minutes and the unbound supernatant was transferred into a new tube and kept for analysis. The immunoprecipitated proteins bound to the beads were washed 5 times with 1XPBS containing 10% glycerol and 0.05% Tween-20 and finally eluted using 20 μl of 0.1M Tricine buffer (pH 3) at RT for 10 min. The eluted samples were neutralized to physiological pH using NaOH, mixed at 1:1 ratio with 2xMAL labeling buffer and subjected to dialysis against 1xMAL labeling buffer. For gel analysis, the eluted sample was mixed with final 1XLaemmli sample buffer containing 25 mM Tris-HCl, pH 6.8, 10% (w/v) SDS, 10% (v/v) glycerol and 0.1% (w/v) bromophenol blue, denatured at 95°C for 5 minutes and subjected to separation on 4-12% Bis-Tris SDS-PAGE using 1xMES buffer at 150V for 1h under non-reducing conditions. The proteins were fixed at RT for 2h on a shaking platform using 40% ethanol and 10% glacial acetic acid. Subsequently, the gel was stained with 1X Flamingo stain in milliQ water for overnight at RT on a shaking platform. Finally, the gel images were acquired using the Pharos scanner (Biorad) with laser excitation at 532 nm. For single protein analysis using the nano-channel device, the eluted samples were subjected to the dual-color labeling protocol with Atto565 maleimide and Atto643 NHS ester.

## Supporting information

CA/OVA tracking

VEGF isoforms

## Author Contributions

S. Ohayon, L. Taib, N. C. Verma, I. Bhattacharya and M. Iarossi performed all nano-channel measurements and analyzed the data, D. Huttner and B. Marom prepared protein samples, A. Meller designed and supervised the research. All authors co-wrote the manuscript.

## Acknowledgements

We acknowledge stimulating discussions with Dr. Avner Adini (Harvard Medical School), Dr. Itai Chowers (Hadassa Medical Center). This project has received funding from the European Research Council (ERC) No. 833399 (NanoProt-ID) and ERC-PoC No. 966824 (OptiPore) both under the European Union’s Horizon 2020 research and innovation programme grant agreements. S.O is supported by the Azrieli fellowship foundation. N.C.V. has been partially supported at the Technion by a fellowship of the Israel Academy of Science and Humanities and the Israel Council for Higher Education.

