## Supporting Information for "Single protein molecules separation, tracking and counting in ultra-thin silicon channels"

Department of Biomedical Engineering
Russell Berrie Nanotechnology Institute
Technion -IIT
Haifa, Israel

**Table of content:**

1. Nano-channel device fabrication, assembly and treatment.
2. Electro-optical setup for single molecule imaging.
3. Dual color labeling of SDS-denatured proteins and Degree of Labeling confirmation.
4. Immunoprecipitation and protein labeling procedure.
5. Data clustering analysis and confirmation.
6. Additional VEGF isoforms spike-in serum, and three cytokines nano-channel analyses.

**SI Figures list:**

1. An overview of chip fabrication steps.
2. A schematic illustration of the dual color electro-optical setup.
3. SDS-PAGE analysis of dualy labelled CA and Ova proteins.
4. Degree of Labeling (DoL) estimation using UV-Vis spectrometry.
5. Estimation of DoL for low absorbance proteins.
6. Recovery of hVEGF isoforms spiked into human serum by immunoprecipitation.
7. Labeled IP samples of recombinant VEGF of different concentrations spiked into serum.
8. Mathematical validation of GMM clustering for CA and OVA nano-channels experiment.
9. Mathematical validation of GMM clustering for the 4 cytokines panel.
10. Violin plots for different mixtures of VEGF isoforms.
11. VEGF isoforms spike and recovery violin plots (4D data).
12. Single protein molecule quantification of a 3-cytokine panel using our method.

**SI Tables list:**

1. Properties of proteins used in this study and their theoretical $M_w$ post complete labeling or post complete Cysteine labeling step.
2. Labeling yields of proteins samples used in this study.

### 1. Nano-channel device fabrication, assembly and treatment

#### Device fabrication and assembly

The fabrication is performed on a specialized double-sided 100 mm wafer (SVM, CA, USA) with a distinctive layer arrangement: $SiN_x/SiO_2/Si$ (50 nm/350 nm/350 µm). The wafer is thoroughly cleaned using organic solvents, followed by 5 minutes of baking on a hot plate at 300°C in order to evaporate any residues or organic debris. Subsequent to the cleaning procedure, AZ1518 photoresist is spin-coated onto the wafer at 4000 RPM to achieve a thickness of ~1.8 µm and again baked on a hotplate for 2 mins at 115°C. The wafer pattern is exposed to UV light with an exposure of 85 $mJ/cm^2$ by MicroWriter ML3 (Durham Magneto Optics, UK). The microchannels are then developed in Novo Developer (2.14% TMAH in water) for one minute and then washed with water. Subsequently, the wafer is etched by reactive ion etching (RIE) with 10 sccm $CF_4$ and 10 sccm $O_2$ (75 W, 0.15 mbar) for 3 min. The resist is then removed, using acetone and isopropanol, and the wafer is dried. Next, 0.5 mm through-ports are exposed as individual squares at the backside of the wafer, aligned to the microchannels along with cutlines. The $SiN_x$ and the underlying $SiO_2$ insulating layer are etched by RIE and BOE (9 min, RT), respectively, and then opened by anisotropic KOH etching (33% KOH, T = 65°C), in a custom-made temperature and flow controlled etch station overnight. The whole process is illustrated in SI Figure 1.

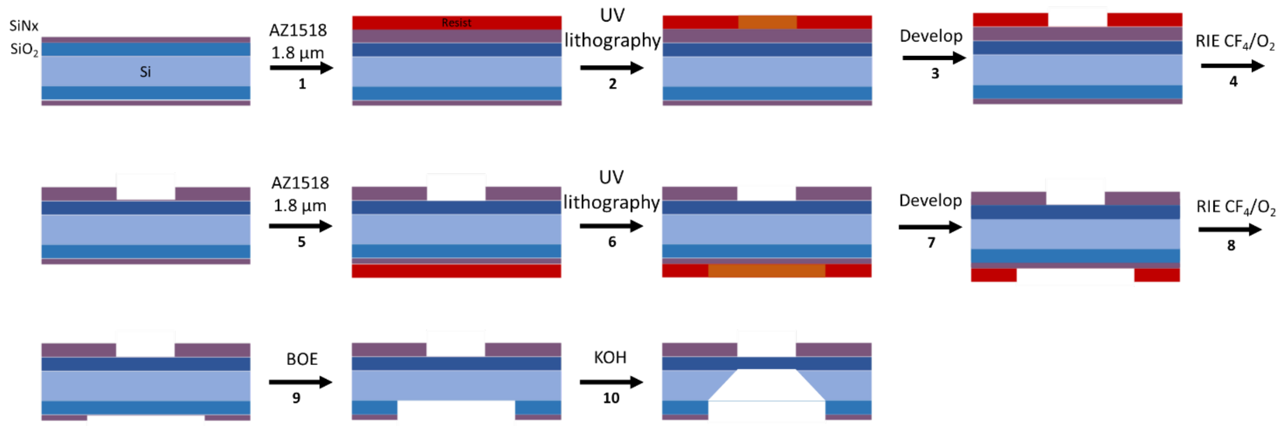


**Figure 1. An overview of chip fabrication steps.** (1) The starting 4-inch wafer consists of different layers: $Si/SiO_2/SiN_x$ (350 µm/500 nm/50 nm) onto which AZ1518 resist is spin-coated to a thickness of 1.8 µm. (2) UV lithography is used to expose the channels structure mask (3) and the channels are developed in Novo Developer. (4) The unprotected pattern is subsequently etched to remove the exposed $SiN_x$ layer by reactive ion etching (RIE) using 1:1 $CF_4$:$O_2$. (5) AZ1518 resist is spin-coated to a thickness of 1.8 µm. (6) Windows to be opened into the channels, and through-ports are backside aligned to the subsequently exposed microchannels using UV; (7) developed in Novo Developer and (8) etched by RIE. (9) The underlying $SiO_2$ is then etched by a buffered oxide etch (BOE) for 9 mins. (10) Silicon is anisotropically etched by 33% KOH overnight.

#### Device assembly and channel coating

Prior to the device assembly, the chips are sonicated to remove the remaining 500 nm of $SiO_2$ films to allow fluid to flush through the silicon substrate. Subsequently, BOE is applied for 3 minutes to the nanochannels side to deepen the channels and slightly clean the SiNx surface. The chips are then cleaned using hot Piranha solution ($H_2O_2$/Sulfuric acid), washed in DDW water, and carefully dried using a nitrogen gun. To facilitate the anodic bonding of glass cover slides with the chips, the front side of each chip (the side where channels are carved) and coverslip are activated using air plasma (0.4 Torr for 5 min). Anodic bonding is performed by placing the glass slide in contact with the wafer at 400°C while simultaneously applying a voltage of 1000 V for roughly 15 minutes. After ensuring the glass is appropriately bonded to the device's front side, the back side (where the ports are carved) is bonded to a ~3 mm thick PDMS slab with pre-cut-through holes aligned with the device's fluid ports.

After the full device is assembled, the interior surfaces of the channels are coated to prevent protein absorption to the surface of the channels and deposition of the SDS-PAGE. In this process, first, the channels are washed with 1 M NaOH for 10 mins, followed by flushing with DDW. Subsequently, the channels are filled with a 2:3:5 mixture of 3-(trimethoxysilyl)propyl methacrylate, glacial acetic acid, and DDW water for 30 mins, then rinsing with methanol and DDW water for 5 mins each. A solution of 5% (w/v) acrylamide coating containing 5 mg/mL 2,2′-azobis(2- methylpropionamide)dihydrochloride (V-50) photoinitiator is flushed through the channels, and the channels are then exposed to 365 nm UV-light in close proximity (< 3 mm) for 10 mins (Spectroline ENB-260C). The channel flushing is done by using positive filtered air pressure of ~ 2 bar.

***In-situ* SDS-PAGE curing in the nano-channels**

The SDS-PAGE used in these experiments was with 8% acrylamide/bis-acrylamide. Gel matrices were prepared by adjusting the total of 40% acrylamide/bis-acrylamide solution (Sigma) with 0.1% (w/v) tris-glycine (0.025 M Tris base and 0.194 M glycine pH 8.3), 0.2% (w/v) SDS and 2,2′-azobis[2-methyl-N-(2-hydroxyethyl)propionamide] (VA-086) photoinitiator. The solution was flushed through the channels, and the chip was taken to a clean room environment for UV exposure. Exposure was done using MicroWriter ML3 by 385 nm wavelength. For curing the gel matrix, 1.5/2.5 mm X 100 µm rectangle patterns were used on the main channel. Both the exposure time and intensity affect the density of the cured gel, which was optimized to yield the best separation results. The UV intensity used was 5000 mJ/ $cm^2$ for all experiments, and time was varied in steps to induce a polymerization gradient.

### 2. An electrooptical setup for dual color single-molecule imaging.

To image the devices, we constructed a custom dual-color single molecule sensing setup that produces uniform illumination of the imaging zone using two laser excitations (532 nm and 640 nm). The lasers exposure times were synchronized with an EM-CCD frame acquisition periods and were alternated to allow independent excitation of “green” (Atto 565) and “red” (Atto 643) fluorophores (Figure 1c). The emitted light was collected using a high NA microscope objective (Olympus 60x/1.45), split to two channels, filtered and imaged by an EM-CCD camera. The nano-channel device was placed on a piezo-driven *Z* axis stage equipped also with long-travel *X-Y* motors, to allow precise placement of the device at the imaging zone as confirmed using a white light image (not shown). To electrokinetically drive the proteins through the nano-channel, a steady voltage was applied using a computer-controlled electrometer. The system was monitored in real-time using a

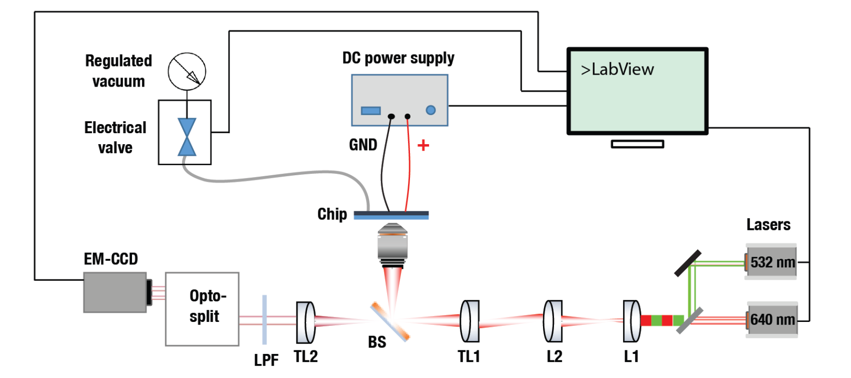


**Figure 2.** A schematic view of custom electro-optical setup to control protein motion in the nano-channel, with alternating lasers excitation for two color single molecule imaging at high speed.

custom LabView program, which in addition to fully control all parts of the system was also used to directly stream the movies to disk from the EM-CCD.

#### 3. Dual color labeling of SDS-denatured proteins and Degree of Labeling confirmation

Proteins (either as individual species or as a mixture of different proteins) were resuspended in Maleimide labeling buffer consisting of 10 mM sodium phosphate, 150 mM NaCl and 1 mM SDS (pH 7.4). Disulfide bonds were reduced using 0.5 mM TCEP at 37°C for 30 minutes and proteins were denatured at 95°C for 5 minutes. After cooling down the solutions at RT for 15 minutes, atto565 maleimide, dissolved in DMSO or buffer, was added to the protein samples at a ratio of 2.6x fold per cysteine residue and incubated at 25°C for overnight with shaking at 300 rpm. Subsequently, the samples were dialysed against 10 mM sodium phosphate, 150 mM NaCl and 1 mM SDS (pH 8.4, adjusted with 0.2 M sodium bicarbonate (pH 9)). Atto643 NHS ester, dissolved in DMSO or in buffer, was added to the protein samples at a ratio of 20x fold per lysine residue and incubated at 25°C for 1h inside a heat block with shaking at 300 rpm. The labeled samples were diluted 10x fold with 10 mM sodium phosphate and 1 mM SDS (pH 8.4) buffer to adjust the NaCl concentration to 15 mM. Unconjugated fluorophores were removed using at least six buffer exchange washes with the protein concentrator VivaSpin of a 30 kDa cutoff. The final volume of the labeled proteins was adjusted with a buffer containing 10 mM sodium phosphate, 15 mM NaCl and 1 mM SDS (pH 8.4). A detailed explanation on how the labeling yield was estimated/calculated is provided in the next section. Qualitative analysis for the labeled proteins was carried out by separating a small fraction of the dually labeled protein samples on a 4-12% Bis-Tris SDS-PAGE and imaged using Pharos scanner (Biorad) with laser excitation at either 532 or 635 nm.

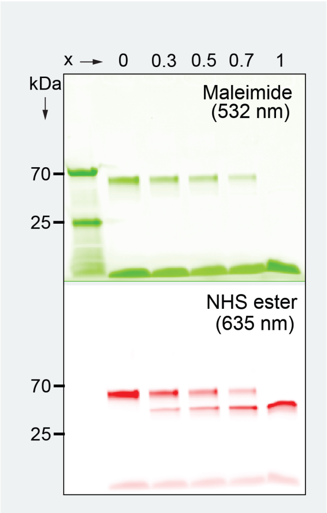


**Figure 3.** Dual-color SDS-PAGE analysis of high-yield covalent bioconjugation showing >90% of K and C residues labeling in Ovalbumin (OVA) and Carbonic Anhydrase (CA). Different molar ratio mixtures of the two proteins with $x = C_{CA}/(C_{CA} + C_{OVA})$ were prepared and co-labelled with the Mal-Atto 565 and NHS-Atto 643 reactive dyes.

**Error! Reference source not found.** consists of the list of proteins used for labeling and nano-channel analysis in this study, as well as their molecular weights before and after completing dual labeling with the two dyes, Atto565 maleimide and Atto643 NHS ester. To determine the proteins' degree of labeling (DoL) we used a combination of dual color PAGE and UV-Vis spectrometry. For proteins that are labeled under native conditions, Atto-Tec datasheet[1] recommends using the following equation for calculating the degree of labeling (DoL), which represents the ratio between the concentration of the dye ($C_{dye}$) to the concentration of the protein ($C_{prot}$) in the labeled sample:

$$Equ\ 1. \qquad DoL = \frac{C_{dye}}{C_{prot}} = \frac{A_{max}\ \varepsilon_{prot}}{A_{prot}\ \varepsilon_{max}} = \frac{A_{max}\ \varepsilon_{prot}}{(A_{280} - A_{max}\ CF_{280})\ \varepsilon_{max}}$$

| Protein | $M_w$ (kDa) | # of C | # of K | $M_w$ post Cys labeling (kDa) | $M_w$ post dual-labeling (kDa) | Additional information |
|---|---|---|---|---|---|---|
| **CA** | 30 | 0 | 18 | 30 | 46.78 | Source: Sigma, cat. # C2273<br>Specie: Bovine (purified from bovine erythrocytes)<br>Used as a marker in SDS-PAGE |
| **OVA** | 44.2 | 6 | 20 | 48 | 66.65 | Source: Sigma, cat. # A7642<br>Specie: Chicken (purified from chicken egg white)<br>Used as a marker in SDS-PAGE |
| **IP-10** | 8.65 | 4 | 10 | 11.18 | 20.51 | Source: ProspecBio, cat. # CHM-330<br>Specie: human (recombinant human IP-10 produced in *E. Coli*)<br>Chemokine of the immune response (belongs to the CXC chemokine family); is elevated in both bacterial and viral infections |
| **TRAIL** | 19.6 | 1 | 11 | 20.23 | 30.49 | Source: ProspecBio, cat. # CYT-546<br>Specie: human (soluble recombinant human TRAIL produced in *E. Coli*)<br>Inflammation response; is used as a biomarker for viral infection |
| **IL-6** | 20.98 | 4 | 14 | 23.51 | 36.57 | Source: Peprotech, cat. # 200-06<br>Specie: human (recombinant human IL-6 produced in *E. Coli*)<br>Immune and inflammation responses |
| **CRP** | 23.2 | 2 | 13 | 24.47 | 36.59 | Source: ProspecBio, cat. # PRO-335<br>Specie: human (recombinant human CRP produced in *E. Coli*)<br>Inflammation response, is used as a biomarker for bacterial infection |
| **$VEGF_{165}$ monomer** | 19.1 | 16 | 11 | 29.23 | 39.49 | Source: Peprotech, cat. # 100-20<br>Specie: human (recombinant human VEGF165 produced in *E. Coli*)<br>Cytokine, promotes angiogenesis and vascular permeability |
| **$VEGF_{121}$ monomer** | 14.06 | 9 | 7 | 19.76 | 26.29 | Source: Peprotech, cat. # 100-20A<br>Specie: human (recombinant human VEGF121 produced in *E. Coli*)<br>Cytokine, promotes angiogenesis and vascular permeability |

**Table 1. Properties of proteins used in this study and their theoretical $M_w$ post complete labeling or post complete Cysteine labeling step.**

$A_{max}$ is the absorption of the dye at its absorption maximum and $A_{prot}$ is the absorption of the protein at 280 nm, while subtracting the contribution of the dye absorbance at this wavelength, determined by the correction factor of this dye at 280 nm ($CF_{280}$). The extinction coefficients of the dye and the protein are $\varepsilon_{max}$ and $\varepsilon_{prot}$, respectively. The labeling efficiency in a particular sample can be then calculated by dividing the obtained DoL by the number of possible labeled residues, or $DoL_{max}$, which exist in a particular protein.

$$Equ\ 2. \qquad Labeling\ yield = \frac{100\ x\ DOL}{DOL_{max}}$$

However, we found that for high labeling density per protein cases, Equ 1. is no longer valid since the absorbance of the dyes at 280 nm masks the absorbance peaks of the labeled proteins. To address this, we developed an alternative approach for calculating the labeling yields of the proteins used in this study. As shown in SI **Error! Reference source not found.**, we use UV-Vis absorbance measurements of labeled proteins at 643 or 565 nm to calculate the concentration of the Atto643 or Atto565, conjugated respectively to either lysine or cysteine residues. For proteins with extinction coefficients > 10,000 $M^{-1}cm^{-1}$, we measured the UV-Vis absorbance at 280 nm of an unlabeled protein sample, which underwent the same labeling procedure but without dye addition, in parallel to the dually labeled proteins. The unlabeled protein concentration was calculated according to Lambert-Beer law using the value of the absorption peak at 280 nm, divided by the appropriate $\varepsilon_{prot}$, as mentioned in SI **Error! Reference source not found.**. Subsequently, the calculated final protein concentration of the unlabeled protein sample, was used to estimate the protein concentrations in the dually labeled sample, allowing us to use Equ. 1 and 2 to determine the labeling efficiencies.

For the proteins with extinction coefficients, lower than 10,000 $M^{-1}cm^{-1}$, this calculation method was not practical, as the protein concentration used resulted in low absorbance measurements. For those proteins, we estimated the labeling efficiencies in the following way. First, we calculated the contribution of absorbance of the Atto643 dye at 565 nm (the atto565 absorbance contribution at 643 nm is negligible). To derive this contribution, we used the absorbance spectra of both dyes, measured at a known concentration. We normalized the Atto643 absorption at 643 nm to 1 ($\varepsilon_{643} = 150{,}000$ $M^{-1}$ $cm^{-1}$). Accordingly, the Atto565 dye absorption at its peak (565 nm) is 0.8 ($\varepsilon_{565} = 120{,}000$ $M^{-1}$ $cm^{-1}$) and at this peak value, the Atto643 dye absorption value is 0.066. Hence, the ratio at 565 nm between the absorption of Atto643 and Atto565 is $\frac{0.066}{0.8} = 8.25\%$. Based on this, we can write the following equations for the normalized absorptions:

$$Equ\ 3. \qquad A_{565} = C_{prot}\ \ (N_{Cys}\ \ 0.8 + N_{Lys}\ \ 0.066)$$

$$Equ\ 4. \qquad A_{643} = C_{prot}\ \ N_{Lys}$$

Where $C_{prot}$ is the unknown concentration of a protein, $N_{Cys}$ and $N_{Lys}$ are the number of cysteine and lysine residues that were labeled per protein, respectively. Combining Eqs. 3 and 4, we get the following relationship that does not depend on the concentration of the proteins at all:

$$Equ\ 5. \qquad N_{Lys} = 0.8\ N_{Cys}/[\frac{A_{565}}{A_{643}} - 0.066]$$

Based on the shift in migration in SDS-PAGE observed for atto565 labeled proteins compared with the migration of unlabeled proteins, as shown in SI Figure 3, we found that in most cases, we observe full maleimide labeling (cysteins). This was evident by the fact that we observe effectively a single band in the gel which also corresponded to the estimated shift in molecular weight associated fully labelled proteins with the Atto565 MAL. According to this, $N_{Cys}$, in most cases (only OVA was an exception) equals to the theoretical number of cysteine residues in the protein. For these cases, we can estimate $N_{Lys}$ using Equ 5. We note that this is an estimation only, however it has not affected our ability to discriminate among proteins in the nano-channel. The labeling efficiencies that we obtained for each protein using this strategy are listed in SI Table 2 and marked with a red asterisk.

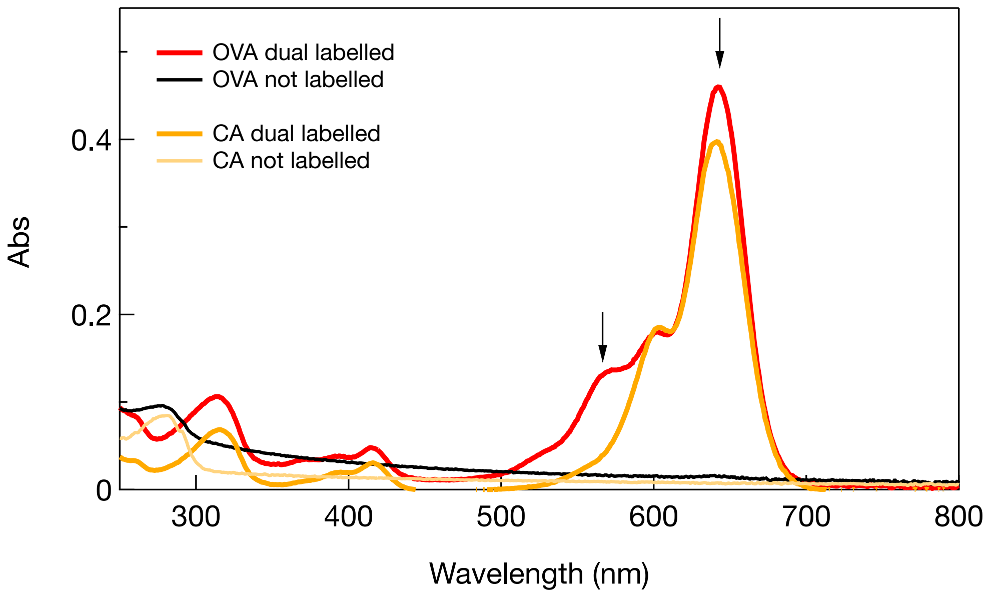


**Figure 4. Degree of Labeling estimation using UV-Vis spectrometry.** Representative absorbance curves of dually labeled proteins are shown for carbonic anhydrase (CA) in yellow and ovalbumin (OVA) in red, as well as the unlabeled CA and OVA samples. The proteins concentrations were estimated based on the 280 nm absorption of the unlabelled proteins (which underwent the same labeling procedure and purification without dye addition) as described. The absorbance spectra of the labelled proteins were scaled by the concentrations to facilitate comparison between the two proteins. The absorbance values at 643 and 565 nm (black arrows) are then used to estimate the dyes cocentrations, using the dyes specific absorption values. Note that the absorption at 565 nm is corrected as follows: $A_{565\_}corrected = A_{565} - A_{643}\ x\ 0.083$. The resulting DoL is given in SI Table 2.

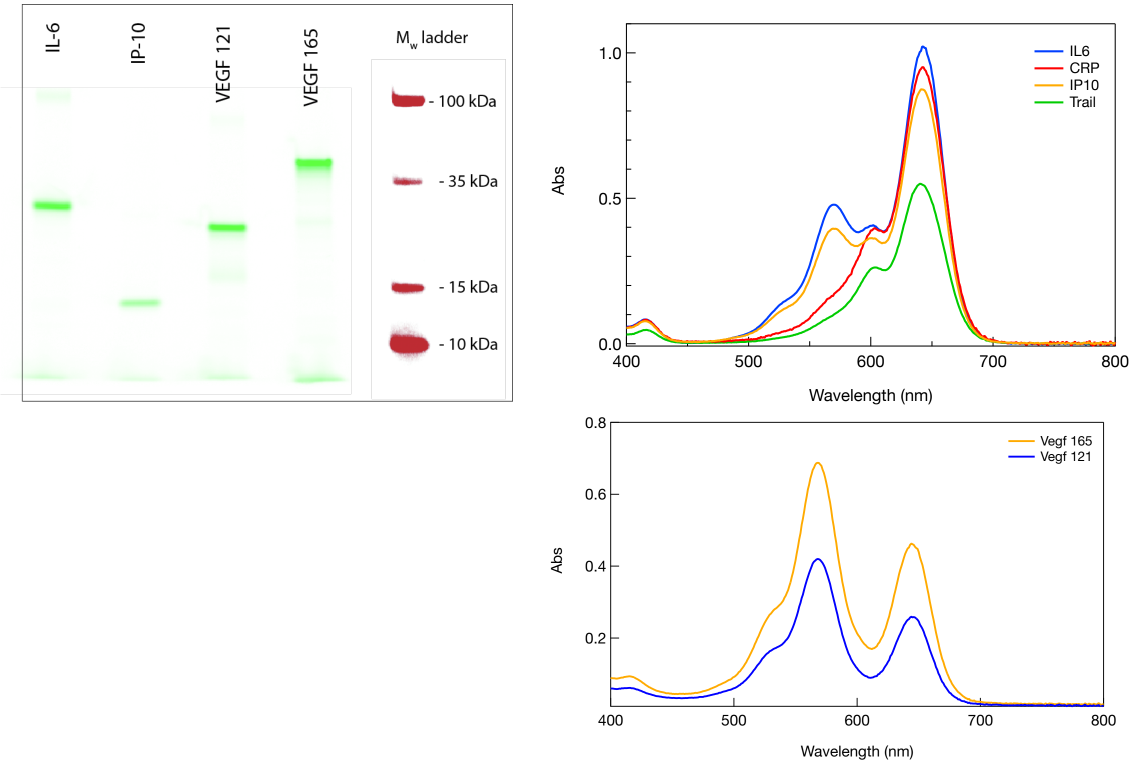


**Figure 5. Estimation of DoL for low absorbance proteins.** Left panel shows an SDS-PAGE scan (532 nm excitation) after Maleimide labeling (Atto-565) only. For the 4 proteins we see clear main single band at molecular weights that correspond to 100% labeling yield. We use the absorbance values at 643 nm and 565 nm, after normalization and Equ. 5 to estimate the DoL of the Lysine residues in each protein. Right panel (top) UV-Vis absorption spectra of the 4 cytokines (IL6, CRP, IP10 and Trail) adjusted for same dilution factor. The relative absorbaence values of thre 4 proteins at both 643 and 565 nm are in line with their expected numbers of K sand C residues, respectively, as indicated in Table 1. Right panel (bottom), UV-Vis spectra for VEGF 165 and VEGF 121 isoforms measured at similar dilutions. Their absorbence levels at 565 and 643 nm are in line with their respective number of C and K residues, respectively.

### 4. Immunoprecipitation and protein labeling procedure

To ensure complete (or near complete) denatured protein labeling, the concentration of either reactive dyes (atto565 maleimide* for cysteine or atto643 NHS ester** for lysine labeling) must be adjusted according to the protein concentration and number of C and K residues per protein. Based on our optimization studies, for protein concentrations > 1 µM, the atto565 maleimide concentration in the reaction is calculated by multiplying the protein concentration times # of cysteines (in protein) times 2.6. This last factor was determined empirically. To avoid dye losses and consequently lower DoL, its concentration in the reaction tube should always be maintained above 2 µM (“basal value”). For NHS ester conjugation, for protein concentrations > 1 µM, we calculate the

| Protein | Extinction coefficient ($M^{-1}cm^{-1}$) | # of C | # of K | Cys DoL | Lys DoL | Method used for DoL calculation |
|---|---|---|---|---|---|---|
| **CA** | 50,420 | 0 | 18 | NA | 100 | Based on absorbance measurements of the dyes and the unlabeled protein sample |
| **OVA** | 31,400 | 6 | 20 | 90 | 100 | Based on absorbance measurements of the dyes and the unlabeled protein sample |
| **IP-10** | 0 | 4 | 10 | 100* | 83 | Based labeling equation |
| **TRAIL** | 27,390 | 1 | 11 | 100 | 77 | Based on absorbance measurements of the dyes and the unlabeled protein sample |
| **IL-6** | 9,970 | 4 | 14 | 100* | 63 | Based labeling equation |
| **CRP** | 44,920 | 2 | 13 | 100 | 100 | Based on absorbance measurements of the dyes and the unlabeled protein sample |
| **$VEGF_{165}$ monomer** | 5,960 | 16 | 11 | 100* | 100 | Based labeling equation |
| **$VEGF_{121}$ monomer** | 5,960 | 9 | 7 | 100* | 100 | Based labeling equation |

Error! Reference source not found. Typical labeling efficiencies of the proteins used in this study. In cases when the extinction coefficient is sufficiently high, we used an absorbance measurement at 280 nm of the unlabeled protein sample to calculate the protein concentration, as explained in SI Figure 2. The unlabeled protein concentration is then used as the estimated concentration of the labeled samples. In cases when the extinction coefficient is lower 10,000 $M^{-1}cm^{-1}$, we estimated the labeling efficiency of the cysteines by observing the migration shift of the atto565 maleimide labeled proteins in the SDS-PAGE, compared to the unlabeled proteins and known molecular markers. In all cases, we obtained a maximal shift and considered the cysteine labeling efficiency to be near full. Then, we used Equ. 5, to calculate the lysines labeling efficiency.

atto643 NHS ester concentration by multiplying the protein concentration times the # of lysines times ~50 fold. Here the basal protein concentration is 1 µM, under which we used a minimal atto643 NHS ester concertation of 200 µM.

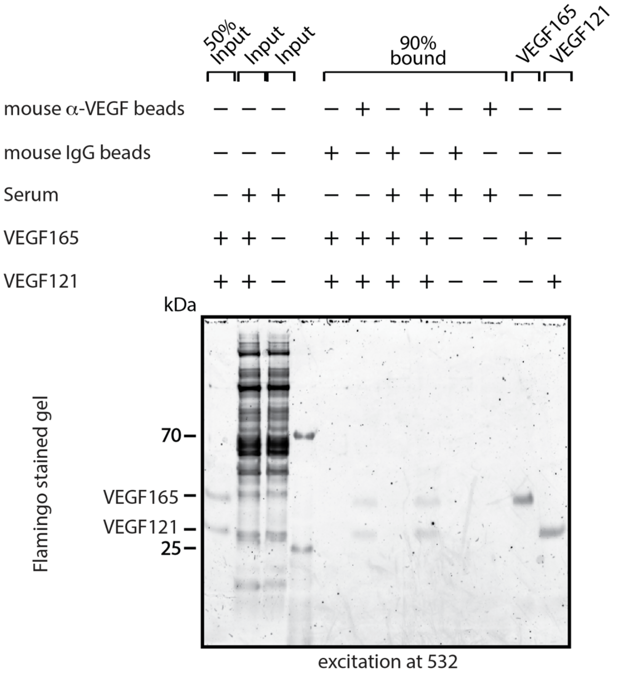


**Figure 6. Recovery of hVEGF isoforms spiked into human serum by immunoprecipitation.** Samples containing either recombinant hVEGF isoforms mixture (VEGF165 + VEGF121 at 1:1 ratio) or human serum either spiked with or without these recombinant VEGF isoforms were subjected to immunoprecipitation with mouse anti-VEGF antibody-conjugated tosylactivated beads for overnight at 4°C on a tube rotator. Tosylactivated beads conjugated to normal mouse IgGκ were used as a negative control. Bound material was washed five times, eluted in 0.1M citrate buffer (pH 3) and mixed at 1:1 ratio with 20 mM sodium phosphate, 300 mM NaCl and 2 mM SDS (pH 7.4). The samples were dialyzed, using mini-dialysis tubes of 3.5 kDa cutoff, against a buffer containing 10 mM sodium phosphate, 150 mM NaCl and 1 mM SDS (pH 7.4). Dialyzed samples were mixed with 1xlaemmli sample buffer containing 25 mM Tris-HCl, pH 6.8, 10% (w/v) SDS, 10% (v/v) glycerol and 0.1% bromophenol blue, denatured at 95°C for 5 minutes and separated on 4-12% Bis-Tris SDS-PAGE using 1XMES buffer under **non-reducing** conditions. The proteins were fixed using 40% ethanol and 10% glacial acetic acid for 2h at RT on a shaking platform and stained overnight with 1X flamingo stain. The gel images were acquired using the Pharos apparatus with the excitation laser at 532 nm. Two clear bands corresponding to the two VEGF isoforms are detected in the lanes containing serum spiked samples which were enriched using the mouse anti-VEGF conjugated beads.

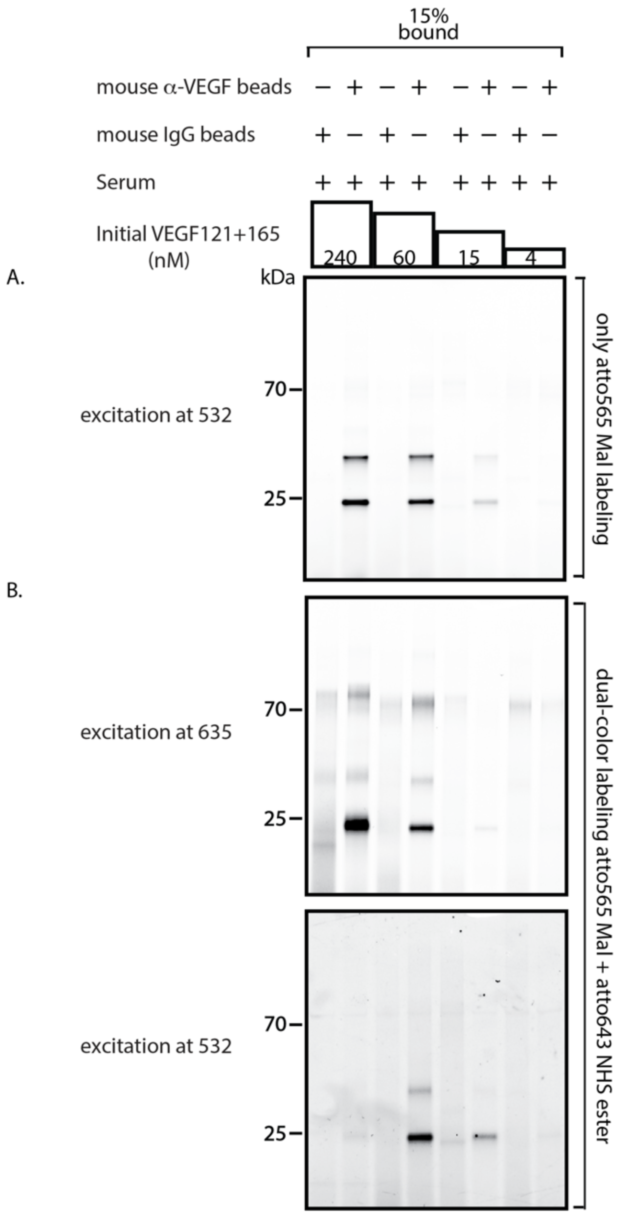


**Figure 7. Labeled IP samples of recombinant VEGF of different concentrations spiked into serum.** Different initial concentrations of recombinant hVEGF isoform mixtures (as annotated above) were used to spike human 7 µl serum samples, before subjecting them to Top14 high abundant protein depletion mini columns. Each sample was subjected to immunoprecipitation according to the procedure described in SI Figure 4 above. Dialyzed samples were reduced with TCEP for 30 minutes at 37°C, denatured at 95°C for 5 minutes and then labeled with atto565 maleimide for O/N at 25°C. These cysteine-labeled samples were dialyzed, using mini-dialysis tubes of 7 kDa cutoff, against a buffer containing 10 mM sodium phosphate, 150 mM NaCl and 1 mM SDS (pH 8.4). At this stage, a small fraction of the samples (15%) was separated on SDS-PAGE and imaged using the Pharos apparatus with the excitation laser at 532 nm (panel A). The atto565 labeled IP samples were further subjected to labeling with atto643 NHS ester for 1 h at 25°C. Finally, unconjugated dyes were removed using buffer exchange washes with a protein concentrator. After purification, a small fraction of the samples was separated on SDS-PAGE and imaged using the Pharos apparatus with the excitation lasers at 532 or 635 nm (panel B). As can be seen in panel A, the two isoforms can be detected in the gel using 532 nm excitation while spiked down to 15 nM concentration. On the contrary, excitation using 635 nm resulted in detection of the two isoform bands only when spiked down 60 nM.

### 5. Data clustering analysis and confirmation

For each nano-channel experiment, we obtain single protein dataset to which we apply the GMM, as explained in Figure 2, to cluster the data based on four parameters, migration time, velocity, green and red emission signals. Often, just by looking at the distribution of these parameters, we can already estimate the minimal number of clusters. Such a case is presented in Figure 2, in which both the migration time and the green emission signal present a clear separation of two clusters. This result is consistent with our design of this experiment, separating dually labeled OVA and CA proteins, and shows the importance of each parameter. However, when dealing with more complicated samples, such as those presented in Figure 5, this visualization approach is insufficient. Although we often a priori know the number of clusters or protein groups we expect to have in a particular sample, we need to mathematically prove that based on the GMM, we have indeed obtained the optimal number of clusters within a dataset. To this end, we used the Calinski – Harbatsz Score[2]. This score is calculated according to the following equation:

$$Equ\ 6. \qquad \text{CH Score} = \frac{\sum_{k=1}^{K} n_k \ ||C - C_k||^2}{\sum_{k=1}^{K} \sum_{i=1}^{n_k} \left|\left|X_{ik} - C_k\right|\right|^2} \frac{N-K}{K-1}$$

$n_k - number\ of\ data\ points\ in\ the\ k^{th}\ cluster$

$C - the\ mean\ value\ of\ the\ clusters\ centers$

$C_k - the\ center\ of\ the\ k^{th} cluster$

$N - number\ of\ all\ the\ data\ points$

$K - the\ number\ of\ clusters$

$X_{ik} - i\ data\ point, in\ cluster\ k$

According to Equ 6., this score shows the ratio between the variance of the cluster's centers, to the variance inside each cluster. In an ideal clustering, the variance between the cluster's centers should be as high as possible and the variance inside each cluster should be as small as possible. Hence, for the optimal number of clusters, we get the maximal CH score values. In this equation, the minimal number of clusters allowed is two. Thus, the score for one cluster is set to be zero. In SI Figure 6, we demonstrate this mathematical validation for the experiment presented in Figure 2. As expected, the maximal CH score is obtained for two clusters, one consisting of CA proteins whereas the other one is consisting of OVA proteins.

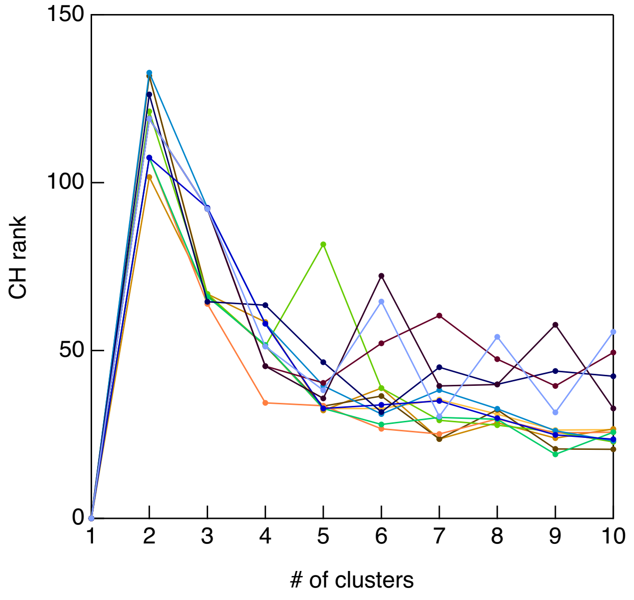


**Figure 8. Mathematical validation of GMM clustering for CA and OVA nano-channels experiment.** The Calinski – Harbatsz score for Figure 2, in which we validate our nano-channel analysis method using CA and OVA proteins, is presented here. The score for this experiment is consistently highest for two clusters, as expected, providing mathematical confidence to our cluster determination and separation.

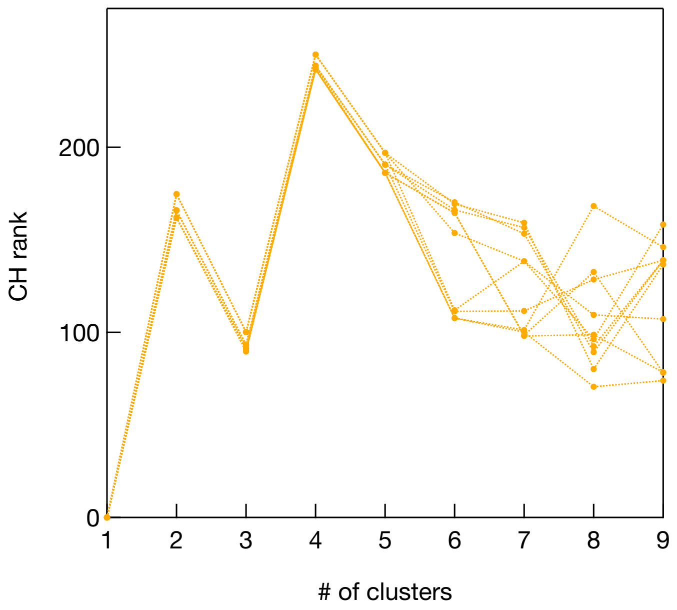


**Figure 9. Mathematical validation of GMM clustering for the 4 cytokines panel.** The Calinski – Harbatsz score for Figure 5. The score for this experiment is consistently highest for cluster of 4.

### 6. Violin plots for the VEGF isoforms discrimination and for 3 cytokines panels

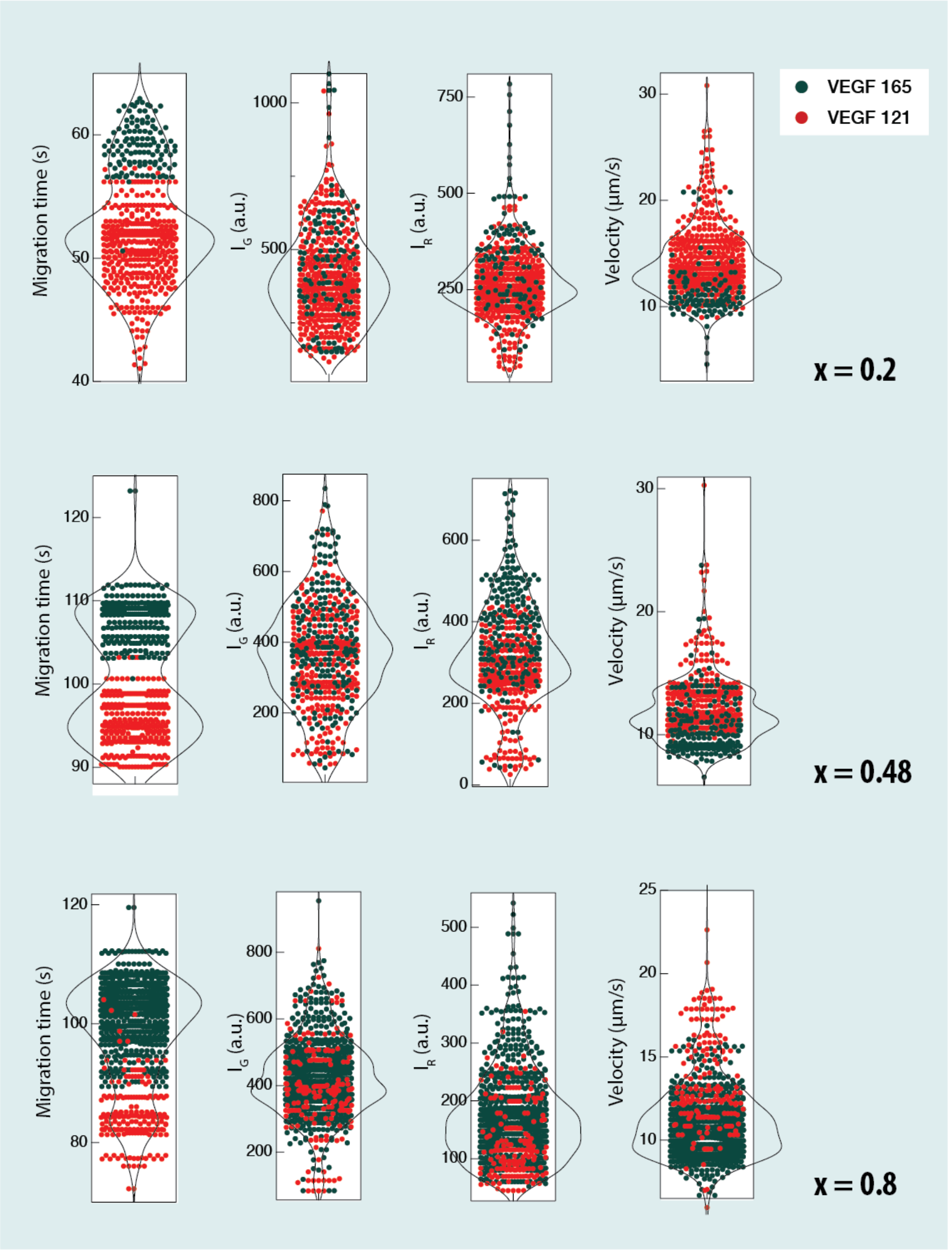


**Figure 10. Violin plots for different mixtures of VEGF isoforms.** Three mixtures of dually labelled VEGF 121 and VEGF 165 with different relative concentrations $x = C_{165}/(C_{165} + C_{121})$ analyzed using our method. The resulting violin plots show distinct two groups with well separated migration times. The faster migration proteins also show lower red laser excited fluorescence consistent with the annotation of this group as the VEGF 121 ($M_w$ = 14 kDa) marked in red, whereas the slower group marked in green is consistent with VEGF 165 ($M_w$ = 19 kDa).

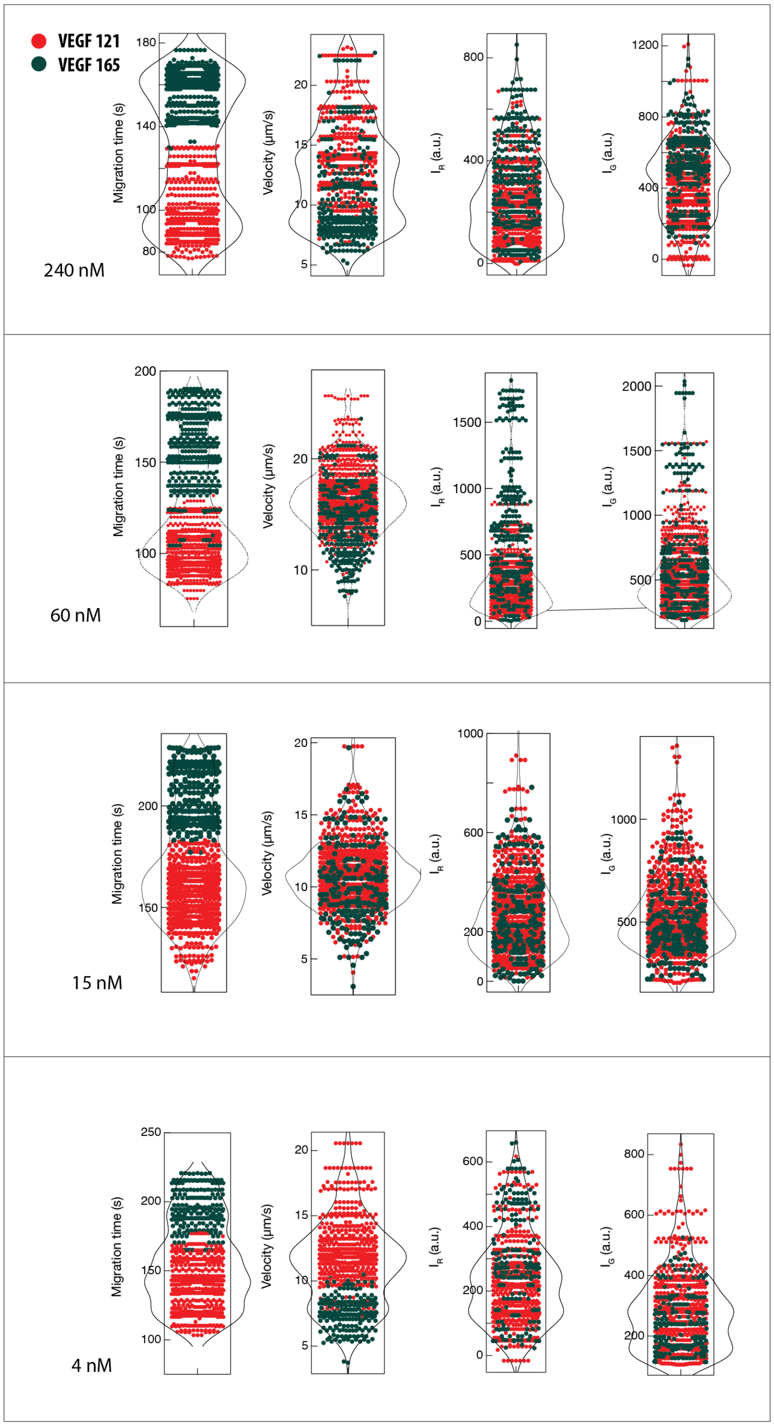


**Figure 11. VEGF isoforms spike and recovery violin plots (4D data).** Human sera was spiked with four total VEGF concentrations both at a VEGF 165 to total VEGF ratio x = 0.3: 240 nM, 60 nM, 15 nM and 4 nM top to bottom panels, respectively. Samples were analyzed after removal of most abundant sera proteins using Top-14 column, pull-down of the VEGFa proteins from spiked sera was performed using custom antibody coated magnetic beads that recognize both isoforms equally. The sample is washed and VEGF isoforms are eluted and undergo dual color labeling (C and K specific), and analysis using the nano-channel device. The samples were significantly ($10^3$ fold or more) diluted before nano-channel analysis. VEGF121 (red) and VEGF 165 (brown) were identified and counted as described in the main text.

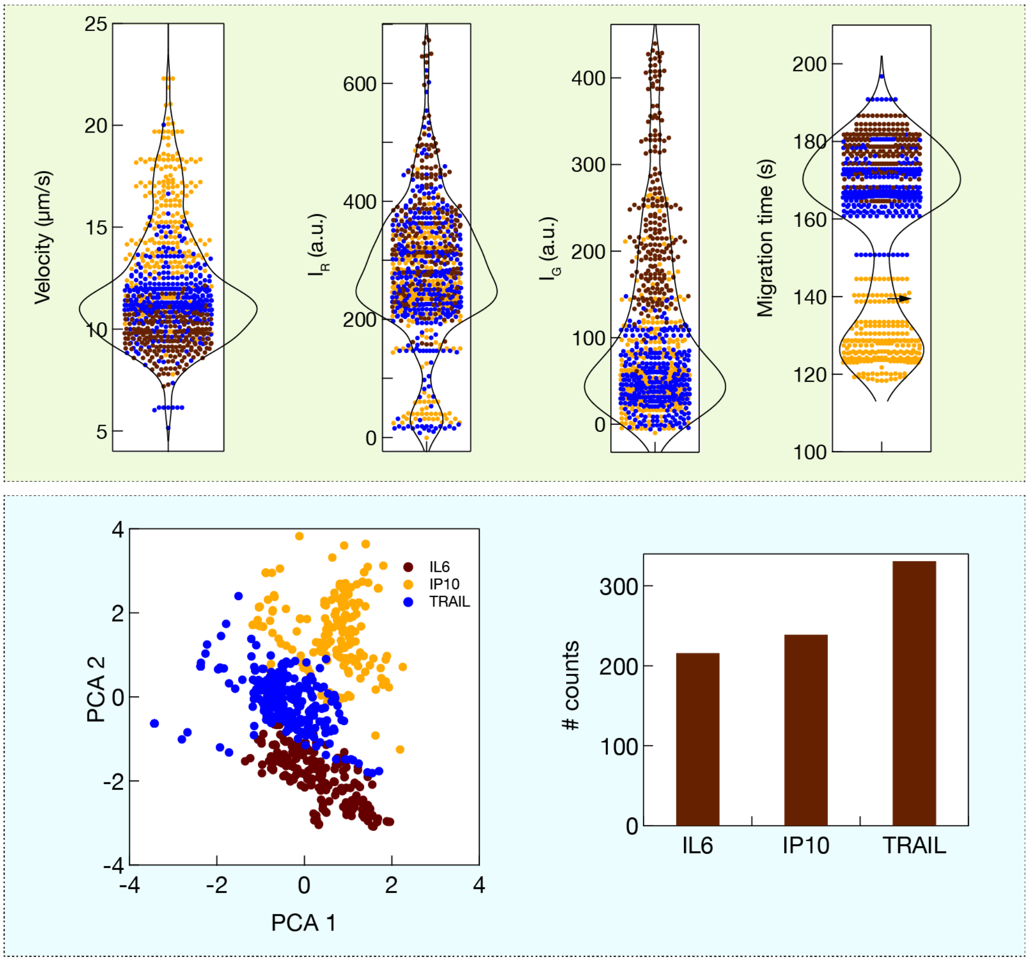


**Figure 12. Single protein molecule quantification of a 3-cytokine panel.** (Top) Violin plots of the 3-cytokine mix run in nano-channels. Despite the similarity in $M_w$ among some of the proteins, our method can distinguish among the 3 cytokines. (Bottom) PCA plot with cytokine annotation based on the GMM analysis is shown in left. The counts of each of the 3 proteins are shown on the right, permitting a direct quantification of the sample.
